# Mapping Gene Expression to an Interpretable Semantic Space

**DOI:** 10.64898/2026.09.11.750957

**Authors:** Xiaoyu Duan, Manu Aggarwal, Vipul Periwal

## Abstract

Cell embeddings organize single-cell expression data, but their dimensions have no biological meaning, so clusters are interpreted afterward. We present MESIC (Mapping Expression to Semantic space with Interpretable Components), which builds the written knowledge about genes held in curated databases into the dimensions themselves. A biomedical language model converts each gene’s summary into a semantic embedding. MESIC compresses these embeddings into a small number of components, each concentrated on a small set of genes and explained by their annotations. The components are computed once from the summaries, so any expression dataset can be mapped onto them, and every cluster, outlier, or cell-type assignment is then characterized by named genes. In cardiomyocytes, outliers in the component space were enriched for hypertrophic cardiomyopathy. In a lung atlas, unsupervised clusters in that space matched the broad cell types that experts had annotated. In both, the components that separated the cells matched their known biology. For about half of the cells that the atlas itself had left unannotated, the same space gave a confident cluster assignment, and with it an interpretation through component-associated genes. Gene summaries thus give single-cell analysis a coordinate system in which every result is traced to genes and what is written about them.

## 1 Introduction

Single-cell RNA sequencing (scRNA-seq) measures the expression of tens of thousands of genes in individual cells. It is used to study how expression profiles vary across tissues, disease states, and perturbations. Standard workflows reduce these high-dimensional, noisy profiles to compact embeddings in which cells can be compared, visualized, and clustered (Satija et al., 2015; Wolf et al., 2018; Luecken and Theis, 2019). More recently, single-cell foundation models have been trained on large collections of expression profiles to produce general-purpose embeddings (Theodoris et al., 2023; Cui et al., 2024; Hao et al., 2024). These embeddings can be reused across datasets and tasks with little or no task-specific training. Cell embeddings from standard workflows and from foundation models alike are not interpretable in themselves. Their dimensions have no direct biological meaning. Clusters are therefore interpreted afterward, through differential expression and pathway analysis. These embeddings also leave out the written knowledge about genes, such as the gene summaries held in curated databases, because they are built from expression profiles alone.

Written gene summaries can be brought into the analysis through pretrained language models, which convert a summary into a semantic gene embedding, a vector that encodes the meaning of its text. GenePT (Chen and Zou, 2025) combined semantic gene embeddings with expression measurements by representing a cell as the average of the semantic embeddings of its genes, weighted by their expression in that cell. On cell-type annotation and other standard tasks, these cell embeddings, built without any pretraining on expression data, matched or exceeded foundation models pretrained on the expression profiles of millions of cells. scELMo (Liu et al., 2026) built on GenePT by having a language model write the descriptions of genes and of cell metadata, rather than taking them from a database.

Like an expression embedding, however, a semantic gene embedding has dimensions with no direct biological meaning. A cell embedding built from it inherits those dimensions, so its clusters must again be interpreted afterward. MESIC (Mapping Expression to Semantic space with Interpretable Components) instead makes the dimensions themselves interpretable. It compresses the semantic gene embeddings into a small number of components, each concentrated on a small set of genes, so a component is defined by those genes and explained by their summaries and annotations (Figure 1A). The components are computed once from the gene summaries, before any expression data are seen, so the same components serve every expression dataset. Mapping single-cell expression profiles onto them places every cell in the same space (Figure 1B). A cluster, an outlier, or a transferred label in this space is then characterized by the components that separate it, and so by named genes. Interpretation is thus built into the dimensions rather than added afterward through differential expression.

**Figure 1:**
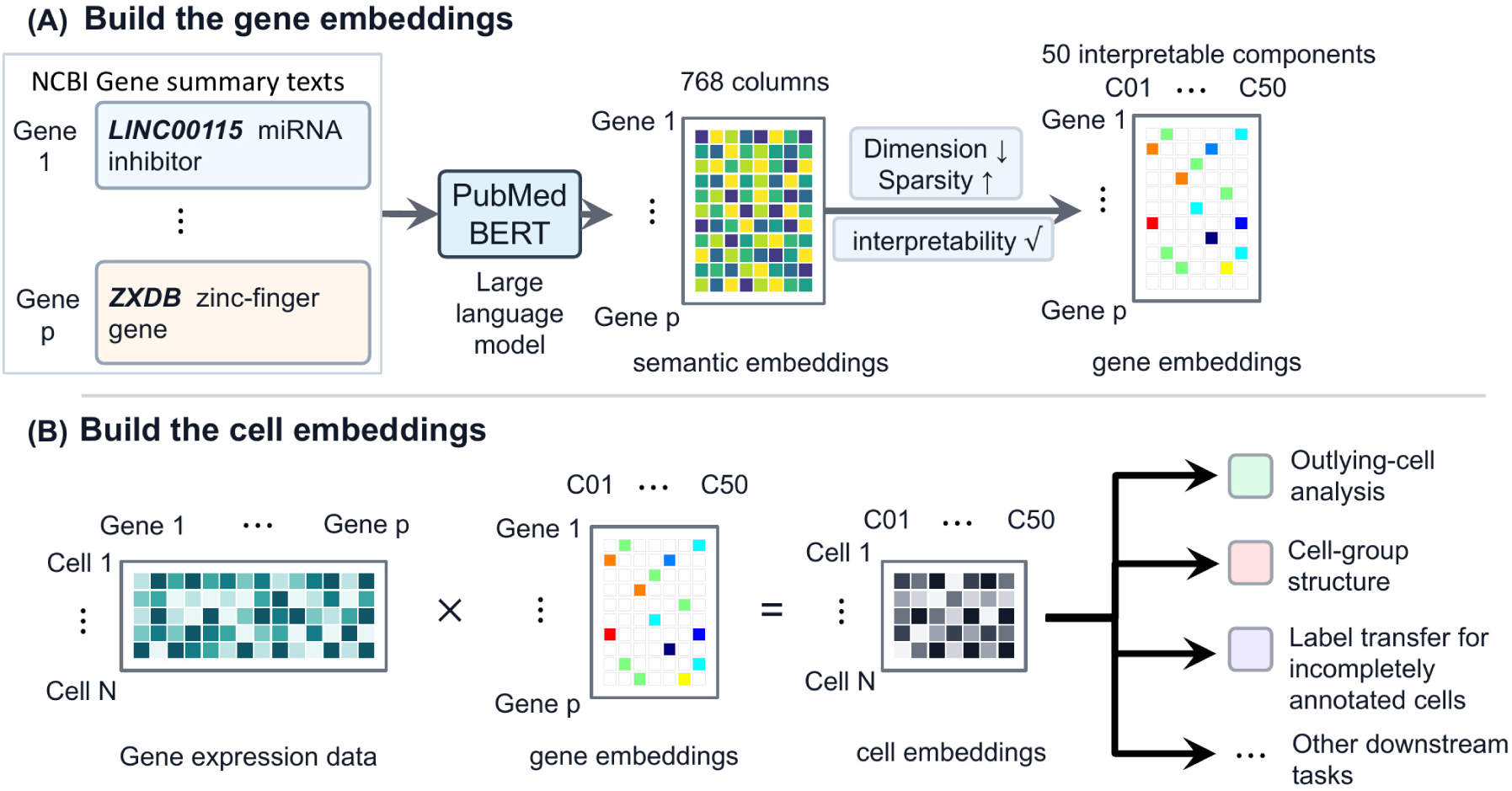
Overview of the workflow. (A) Gene summary texts from the National Center for Biotechnology Information (NCBI) are embedded with PubMedBERT to obtain semantic gene embeddings for 21,788 genes. These semantic embeddings are compressed into a 50-component MESIC gene embedding, with components denoted *C*_1_, …, *C*_50_. Each component is interpreted from its dominant genes and the biological annotations of those genes. (B) Single-cell gene expression data are mapped onto the MESIC gene embedding to obtain MESIC cell embeddings. These cell embeddings support downstream analyses of outlying cells, cell-group structure, incompletely annotated cells, and other tasks.

We test MESIC in two stages. First we test the components themselves, the compressed gene embedding, asking whether the compression loses gene-type information and whether individual components correspond to known biological themes (Section 2.1). Second we test cell embeddings, obtained by mapping two expression datasets onto the same fixed components. For a cardiomyopathy dataset, we rank cardiomyocytes by how far each lies from the others in the component space. We test whether the most distant cells are enriched for disease and identify the components that set those cells apart (Section 2.2). For the Human Lung Cell Atlas, we cluster the cells of one contributing dataset without using their annotations. We compare the clusters with the atlas annotation and identify the components that separate the clusters (Section 2.3). We then use the clusters as a reference for placing atlas cells that the original annotation workflow left incompletely labeled (Section 2.4).

## 2 Results

### 2.1 MESIC preserves gene-type information and yields interpretable components

We used PubMedBERT (Gu et al., 2021), a biomedical language model, to embed National Center for Biotechnology Information (NCBI) gene summary texts for 21,788 genes (Brown et al., 2015). Following the GenePT benchmark (Chen and Zou, 2025), we trained logistic-regression, random-forest, and XGBoost classifiers (Chen and Guestrin, 2016) to predict GENCODE gene-type classes (Frankish et al., 2023) from these semantic embeddings. The classifiers achieved held-out accuracy comparable to that reported in GenePT, indicating that the semantic embeddings captured broad gene-type information.

MESIC compressed these semantic embeddings into 50 components, giving a 50-dimensional MESIC gene embedding for each gene. Gene-type prediction accuracy from these gene embeddings was above 0.96 across all three classifiers (Table 1), suggesting that MESIC’s compression preserved the gene-type information.

**Table 1:**
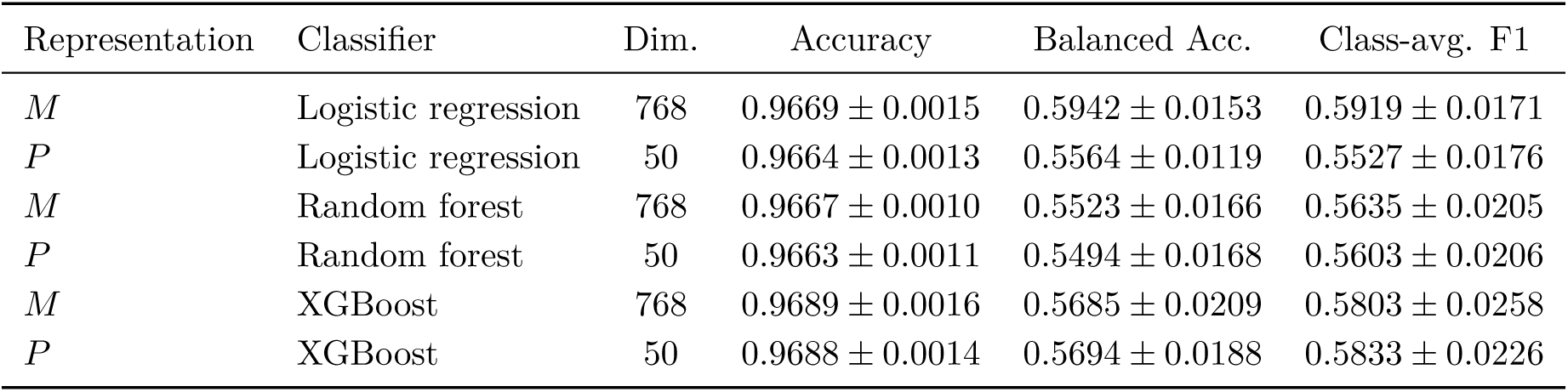
Gene-type prediction using the PubMedBERT semantic embeddings and the MESIC gene embeddings. Values are mean *±* standard deviation across 25 stratified held-out folds. Class-averaged F1 gives equal weight to each gene-type class.

| Representation | Classifier | Dim. | Accuracy | Balanced Acc. | Class-avg. F1 |
| --- | --- | --- | --- | --- | --- |
| *M* | Logistic regression | 768 | $0.9669 \pm 0.0015$ | $0.5942 \pm 0.0153$ | $0.5919 \pm 0.0171$ |
| *P* | Logistic regression | 50 | $0.9664 \pm 0.0013$ | $0.5564 \pm 0.0119$ | $0.5527 \pm 0.0176$ |
| *M* | Random forest | 768 | $0.9667 \pm 0.0010$ | $0.5523 \pm 0.0166$ | $0.5635 \pm 0.0205$ |
| *P* | Random forest | 50 | $0.9663 \pm 0.0011$ | $0.5494 \pm 0.0168$ | $0.5603 \pm 0.0206$ |
| *M* | XGBoost | 768 | $0.9689 \pm 0.0016$ | $0.5685 \pm 0.0209$ | $0.5803 \pm 0.0258$ |
| *P* | XGBoost | 50 | $0.9688 \pm 0.0014$ | $0.5694 \pm 0.0188$ | $0.5833 \pm 0.0226$ |

For both the original semantic embeddings and the MESIC gene embeddings, we summarized the classifier results with row-normalized confusion matrices, in which each row is the annotated gene type and each column is the predicted gene type (Figure 2A,B). Comparing these matrices showed that the two embeddings produced similar patterns of correct and incorrect predictions. Many wrong predictions occurred between gene types whose genes had identical source summaries, making those classes difficult to separate from text-derived information alone. For example, 47 of the 60 TR V genes were predicted as TR J genes. Those 47 genes share their summary text with 60 of the 66 TR J genes. A similar pattern appeared for pseudogenes and protein-coding genes, where 316 pseudogenes shared summaries with 393 protein-coding genes.

**Figure 2:**
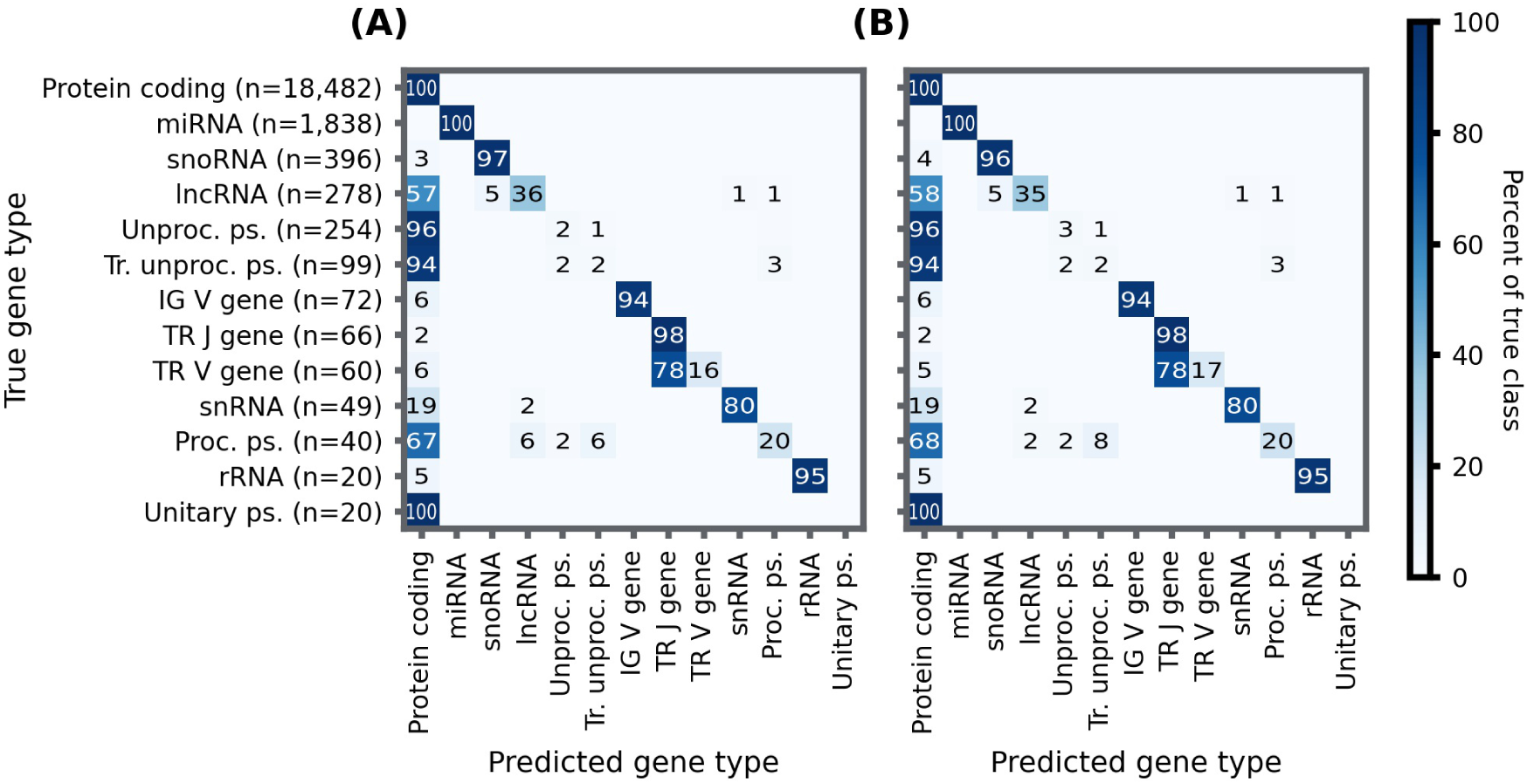
Gene-type prediction from semantic embeddings and the MESIC gene embedding. Row-normalized confusion matrices compare held-out predictions from the 768-dimensional PubMedBERT semantic embeddings (A) and the 50-component MESIC gene embedding (B). Rows denote true GENCODE gene-type classes and columns denote predicted classes. Numbers in row labels give the number of genes in each true class. Matrix entries are pooled held-out row percentages across repeated cross-validation, so each row sums to 100%. Zero-valued entries are left blank. In tick labels, “Proc.” denotes processed, “Unproc.” denotes unprocessed, “Tr.” denotes transcribed, and “ps.” denotes pseudogene.

MESIC was designed so that each component is concentrated on a small set of genes, which we refer to as core genes (Section 3.2 and Supplementary Figure S1). To support component interpretation, we tested the core genes of each component for enrichment against Gene Ontology terms (Ashburner et al., 2000; The Gene Ontology Consortium, 2026), Reactome pathways (Ragueneau et al., 2026), and RNAcentral RNA classes (RNAcentral Consortium, 2021), grouping related enriched terms into themes (Section 3.4). Forty-two of 50 components had at least one supported annotation theme passing the prespecified retention criteria (largest retained adjusted *q* = 0.047). These themes covered diverse biological processes and gene classes, including immune response, ion transport, cilium/flagellum biology, mitochondrial respiration, chromatin regulation, and small-RNA genes (Figure 3).

**Figure 3:**
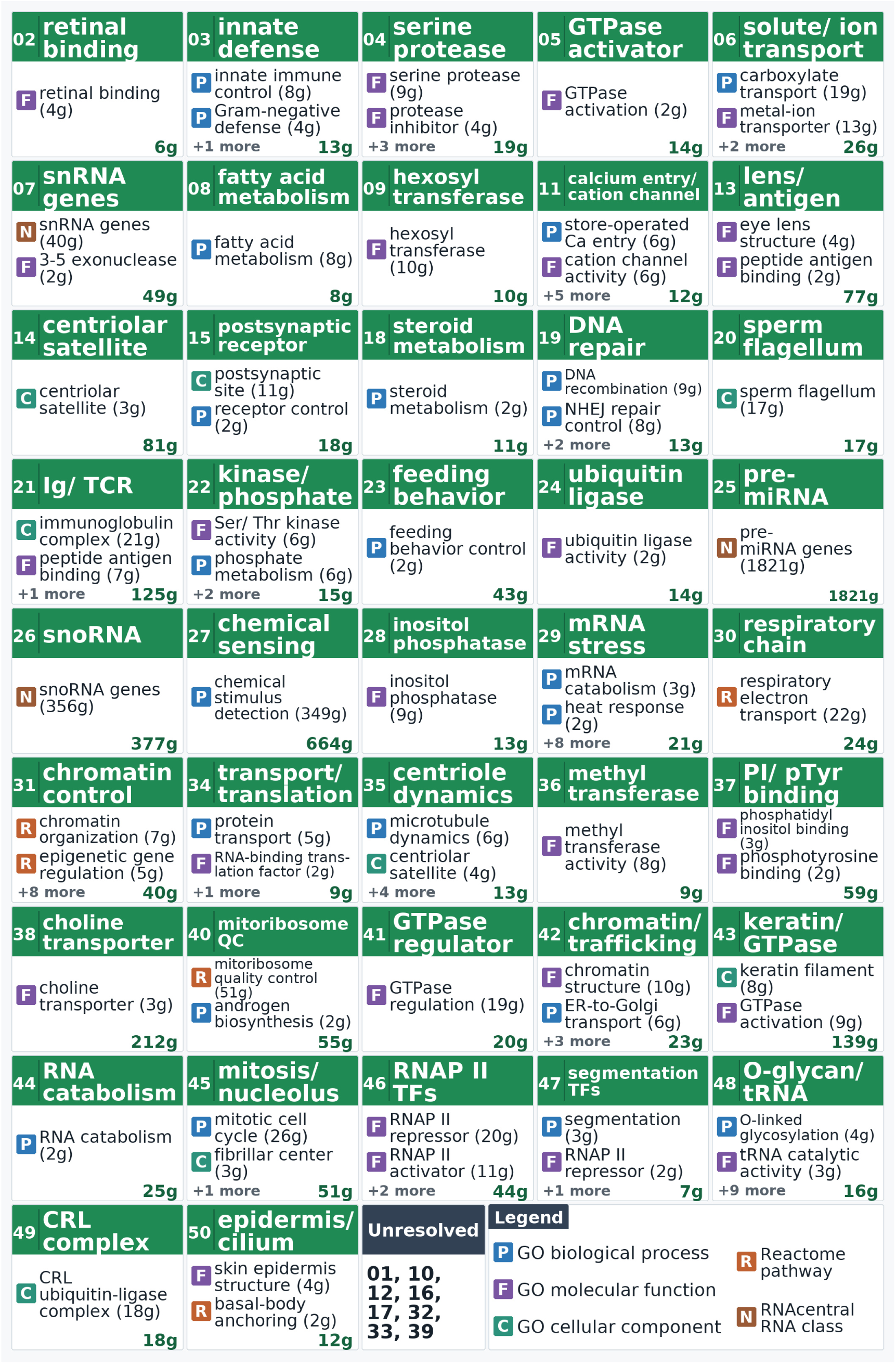
Functional enrichment summaries for the 42 MESIC components with supported annotation themes. Each card shows the component index, a compact biological label derived from retained enrichment themes, and the total number of core genes at the bottom right. Numbers in parentheses after displayed theme labels indicate how many core genes support that theme. A core gene may support more than one retained theme, so theme gene counts within a component are not additive. “+N more” indicates additional retained themes not shown on the card. The unresolved card lists components without supported Gene Ontology, Reactome, or RNAcentral themes. See Supplementary Material for the table listing each component, retained theme, core gene, and the corresponding NCBI summary text.

### 2.2 MESIC cell embeddings link disease-enriched cardiomyocyte outliers to interpretable components

We computed MESIC cell embeddings for 15,846 cardiomyocytes from a human dilated and hypertrophic cardiomyopathy (HCM) single-nucleus RNA-seq dataset (Chaffin et al., 2022). We ranked cells by a multivariate statistical distance from the cardiomyocyte population mean, computed jointly over the components in the MESIC cell embeddings (Section 3.6). The top 1% by this ranking were treated as outliers and were enriched for HCM: 65.4% came from HCM samples, compared with 37.0% of all cardiomyocytes (1.77-fold enrichment; Fisher exact test *p* = 2.6 *×* 10*^−^*^13^; Figure 4A).

**Figure 4:**
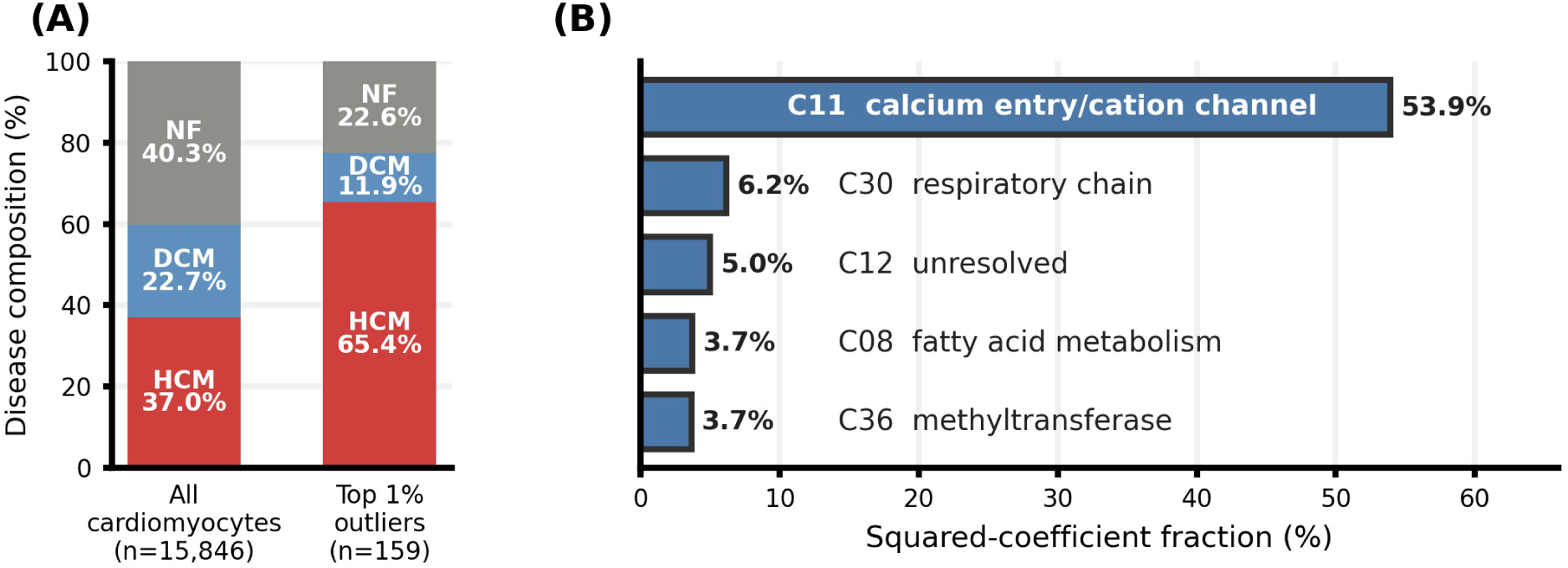
HCM-enriched cardiomyocyte outliers and associated MESIC components. (A) Disease composition of all cardiomyocytes and of the top 1% outliers. (B) Components with the largest squared-coefficient fractions from the sparse logistic-regression model distinguishing the top 1% outliers from the remaining cardiomyocytes.

We then asked which components separate the outliers from the remaining cells. We fit a sparse logistic-regression model that predicts outlier status from the component values, so that the model relies on few components (Section 3.6). To rank the components by how much the model relies on each, we used the squared-coefficient fraction, the fraction of the model’s total squared coefficient weight that falls on each component. Figure 4B shows the five components with the largest fractions. C11 (calcium entry/cation channel) had the largest fraction, consistent with ion-channel and calcium-handling remodeling reported in HCM (Coppini et al., 2013; Helms et al., 2016). C30 (respiratory chain) and C08 (fatty acid metabolism) also had large fractions and were consistent with mitochondrial, energetic, and fatty-acid-oxidation processes reported in human HCM (Nollet et al., 2023; Ranjbarvaziri et al., 2021; Previs et al., 2022). C12 has no supported theme, and we did not find citation-supported interpretations for C36 (methyltransferase) in relation to HCM or abnormal cardiomyocyte biology.

### 2.3 MESIC cell embeddings separate annotation-aligned compartments in a human lung cell atlas

We next computed MESIC cell embeddings for 121,894 cells from the Banovich/Kropski 2020 subset (Habermann et al., 2020) of the Human Lung Cell Atlas (HLCA) core atlas (Sikkema et al., 2023). We refer to this dataset as the Banovich subset below.

We applied unsupervised Leiden clustering (Traag et al., 2019) to these embeddings and selected the final consensus by asking whether repeated random-seed runs returned the same number of clusters and grouped the same cells together. The selected consensus contained six clusters (Figure 5A), with high cluster-count stability and the strongest cell-assignment stability among candidate settings in the reported sweep: 95.3% of seed pairs reached adjusted Rand index (ARI) *≥* 0.95. A separate *k* = 6 spectral clustering (Ng et al., 2002) on the same cell embeddings gave a similar large-scale partition (Figure 5B), supporting that the structure was not specific to Leiden. We refer to the six robust Leiden clusters as L1–L6, with cells that were not consistently assigned to one cluster across runs labeled as uncertain. Full clustering-stability diagnostics are provided in the Supplementary Material.

**Figure 5:**
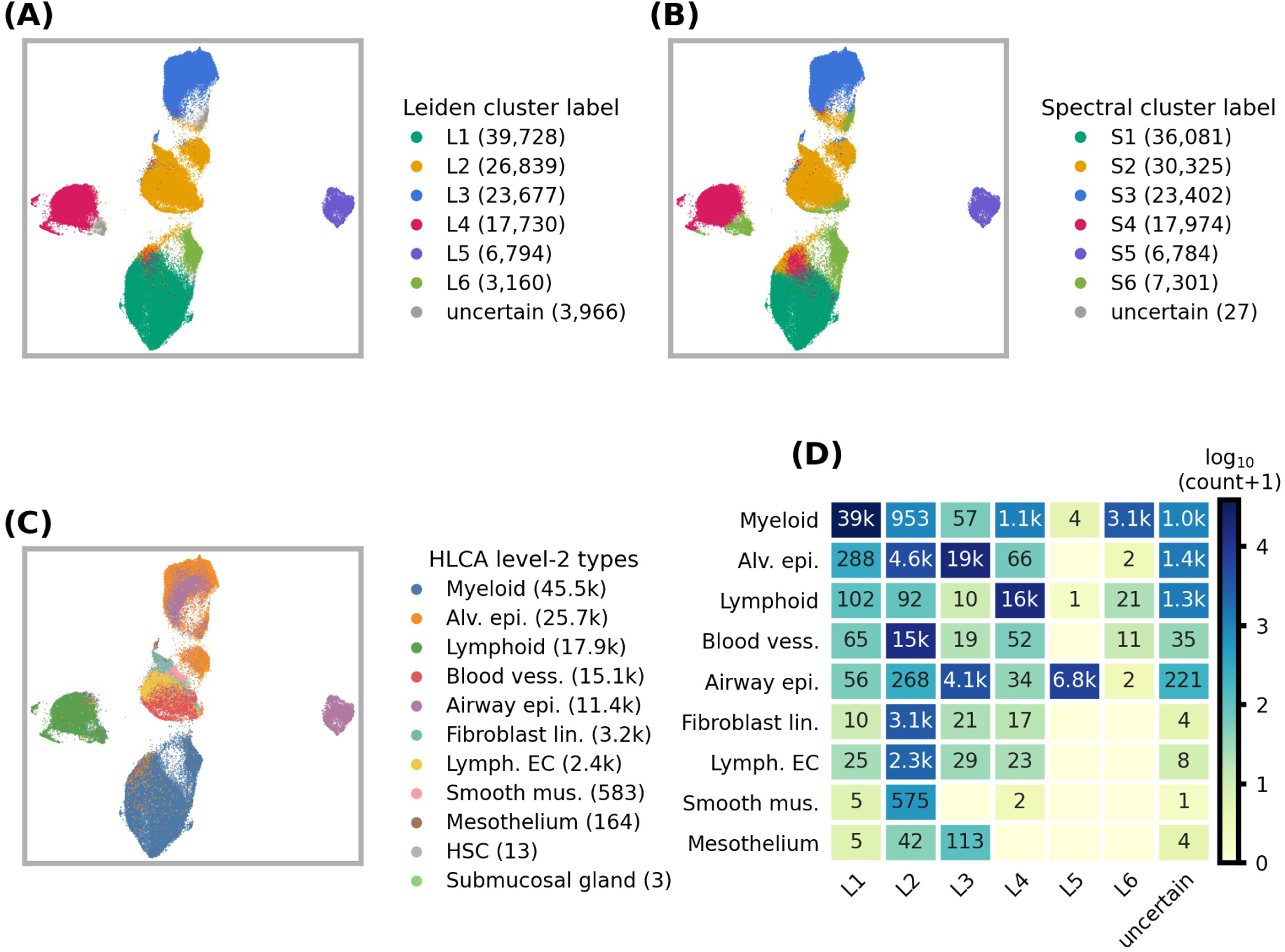
Comparison of MESIC cell-embedding clusters with HLCA level-2 labels in the Banovich subset. (A–C) UMAP visualization of the same MESIC cell embeddings colored by robust Leiden clusters, spectral clusters, and HLCA level-2 labels, respectively. Numbers in parentheses indicate the number of cells assigned to each legend category. (D) Cell-count overlap between HLCA level-2 labels and robust Leiden clusters. Heatmap colors use a log scale, and overlaid text gives raw cell counts. Hematopoietic stem cells and Submucosal Gland cells are omitted from panel D because they contain too few cells. In panels C and D, Alv. epi., Blood vess., Airway epi., Fibroblast lin., Lymph. EC, Smooth mus., and HSC denote alveolar epithelium, blood vessels, airway epithelium, fibroblast lineage, lymphatic endothelial cells, smooth muscle, and hematopoietic stem cells, respectively.

The HLCA core atlas provides expert-curated annotations at multiple levels, from broad ann_level_1 labels to fine ann_level_5 subtype labels. Among these levels, ann_level_2 showed the strongest agreement with the Leiden clusters (ARI = 0.714; Table 2). We therefore refer to ann_level_2 labels as level-2 labels in the remaining HLCA analyses. Figure 5C shows the UMAP colored by level-2 label, and Figure 5D shows the cell-count overlap between level-2 labels and Leiden clusters. Because level-2 labels differed greatly in total cell count, we interpreted this overlap row-wise, asking where cells from each level-2 label were assigned. Lymphoid cells mapped almost entirely to L4, while blood-vessel, lymphatic-endothelial, fibroblast-lineage, and smooth-muscle cells each mapped almost entirely to L2 (*>* 95% for each label). The less concentrated labels showed broader but still structured assignments: alveolar epithelial cells mapped mainly to L3 with a smaller L2 group, airway epithelial cells split mainly between L5 and L3, and myeloid cells mapped mainly to L1 with a smaller L6 group. Thus, L2 was mixed in absolute composition because it collected several smaller stromal and endothelial labels, even though each of those labels was assigned to L2 with high concentration.

**Table 2:** Agreement between high-confidence MESIC Leiden consensus clusters and HLCA annotation levels. Cells labeled uncertain by the Leiden consensus threshold of 0.95 were excluded. For the primary ARI, AMI, and V-measure columns, the same 117,928 cells were used for every annotation level and missing annotation entries were treated as an explicit blank category. The nonblank-only ARI is shown as a diagnostic because finer HLCA levels have incomplete coverage.

| HLCA level | Cells | Coverage | Labels | ARI | AMI | V-measure | ARI nonblank |
| --- | --- | --- | --- | --- | --- | --- | --- |
| Level 1 | 117,928 | 100.0% | 4 | 0.513 | 0.616 | 0.616 | 0.513 |
| Level 2 | 117,928 | 100.0% | 11 | 0.714 | 0.721 | 0.721 | 0.714 |
| Level 3 | 117,928 | 99.4% | 25 | 0.561 | 0.667 | 0.667 | 0.566 |
| Level 4 | 117,928 | 73.0% | 38 | 0.353 | 0.568 | 0.568 | 0.348 |
| Level 5 | 117,928 | 21.5% | 15 | 0.060 | 0.298 | 0.298 | 0.761 |

We then asked which components separated the six clusters and whether those components were consistent with the level-2 labels concentrated in or associated with each cluster. We fit a sparse multinomial logistic-regression model that predicts cluster membership from the component values and ranked the components for each cluster by the same squared-coefficient fraction as in the cardiomyocyte analysis (Section 3.9). Figure 6 shows the five highest-fraction components for each cluster. Several high-fraction components were consistent with broad cluster annotations and prior literature. C21 (Ig/TCR) had a large fraction for the lymphoid-enriched L4 cluster, consistent with lymphocyte antigen-recognition biology (Pishesha et al., 2022). C03 (innate defense) had large fractions for the myeloid-enriched L1 and L6 clusters, consistent with innate immune functions of macrophage and monocyte populations (Hussell and Bell, 2014). C20 (sperm flagellum) had the highest fraction for airway-enriched L5, where the relevant biology is the axonemal machinery shared by sperm flagella and motile airway cilia (Brooks and Wallingford, 2014). C40 (mitoribosome quality control) also had a large fraction for L6 and was compatible with reported links between mitochondrial translation, metabolic state, and macrophage or monocyte activation (Cortés et al., 2023). For L3, the high-fraction components suggested epithelial-relevant themes, including neurotransmitter-receptor signaling, centriole biology, and Rho-family GTPase regulation of epithelial junctions and polarity (Xiang et al., 2007; Nanjundappa et al., 2019; Wallace et al., 2010; Wan et al., 2013).

**Figure 6:**
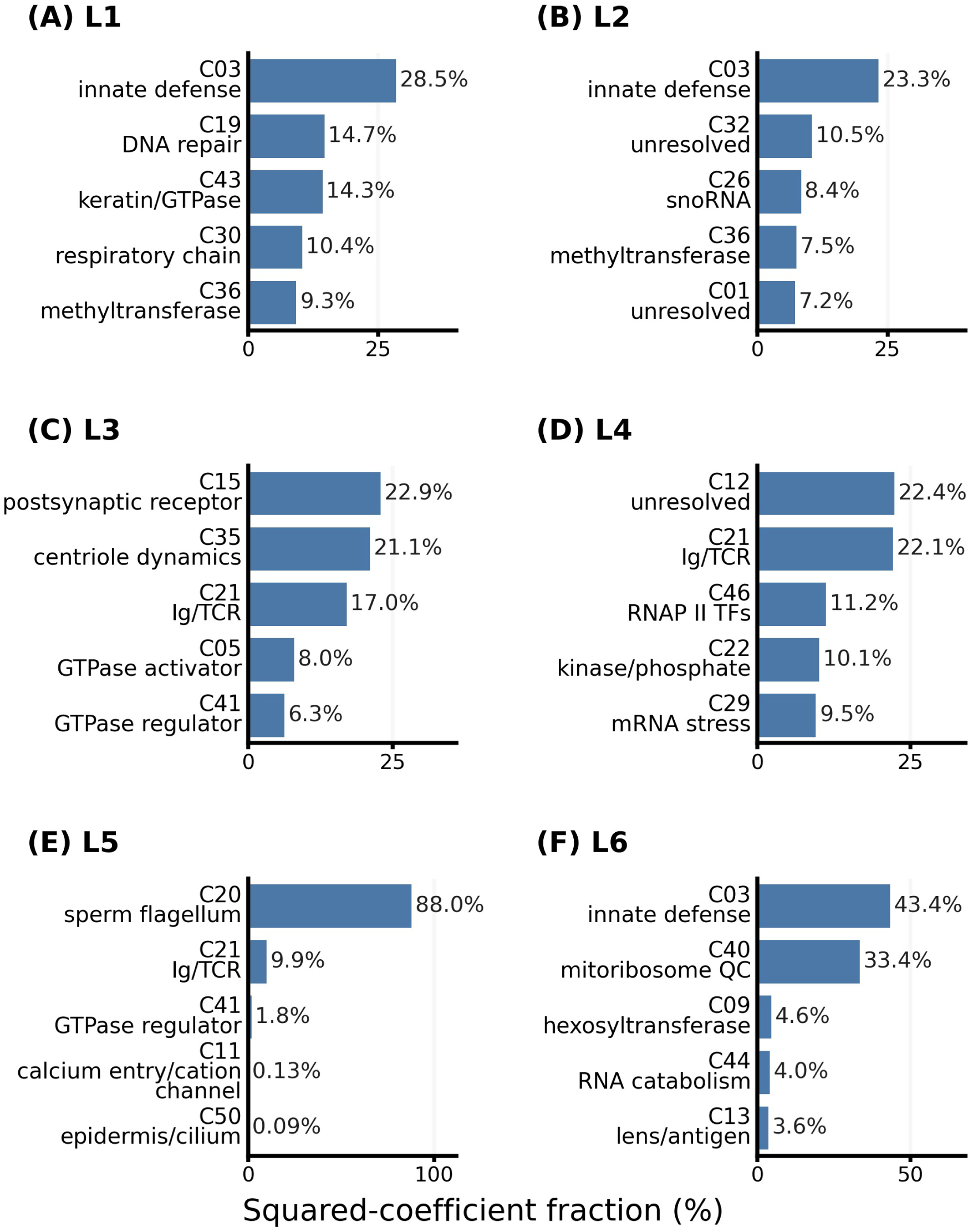
Components distinguishing Leiden clusters in the Banovich subset of the HLCA. For each Leiden cluster, the plot shows the five components with the largest squared-coefficient fractions from the sparse multinomial logistic-regression model.

### 2.4 MESIC cell embeddings provide auxiliary placements for incompletely annotated HLCA cells

The full HLCA contains a core atlas, which includes the Banovich subset, and an extended atlas. In the HLCA annotation workflow, level-2 labels were manually curated for core-atlas cells and then assigned to extended-atlas cells by expression-space transfer from nearby core-atlas cells (Sikkema et al., 2023). This original full-core transfer left 53,331 extended-atlas cells with level-2 label Unknown, meaning that no transferred label reached confidence greater than 0.8. We used these cells as query cells below.

We first repeated this expression-space transfer with one change: the reference was restricted to the Banovich subset, so that both transfers below use the same reference cells. This Banovich-reference transfer assigned confident level-2 labels to 19,044 query cells (36%) under the same confidence threshold of 0.8.

We next mapped the same query cells into the MESIC cell embeddings and transferred the Banovich Leiden labels L1–L6, together with the uncertain consensus label. MESIC confidently placed 28,114 query cells (53%) into one of the six reference clusters. The accepted cells only partly overlapped between the two transfers: 18,468 cells were placed only by MESIC, whereas 9,398 cells were labeled only by the Banovich-reference expression-space transfer. Thus, MESIC provided cluster-level placements for many query cells that did not receive a confident level-2 label from the Banovich-reference expression-space transfer.

The MESIC-only placements were concentrated in a small number of reference clusters (Figure 7A). Most mapped to L2, with a smaller group mapped to L3. Based on the Banovich reference, an L2 placement should not be interpreted as a single level-2 cell type. Instead, L2 was the cluster to which most blood-vessel, lymphatic-endothelial, fibroblast-lineage, and smooth-muscle cells were assigned, while also containing a smaller alveolar epithelial group. L3 was mainly associated with alveolar epithelial cells and a subset of airway epithelial cells. Cells accepted only by the Banovich-reference expression-space transfer were mainly assigned to alveolar epithelium, myeloid, or lymphoid labels (Figure 7B). Together, these results show that MESIC can provide auxiliary cluster-level placements for some cells left unresolved by the original full-core HLCA transfer, while cells with low confidence in both Banovich-reference transfers remain unresolved.

**Figure 7:**
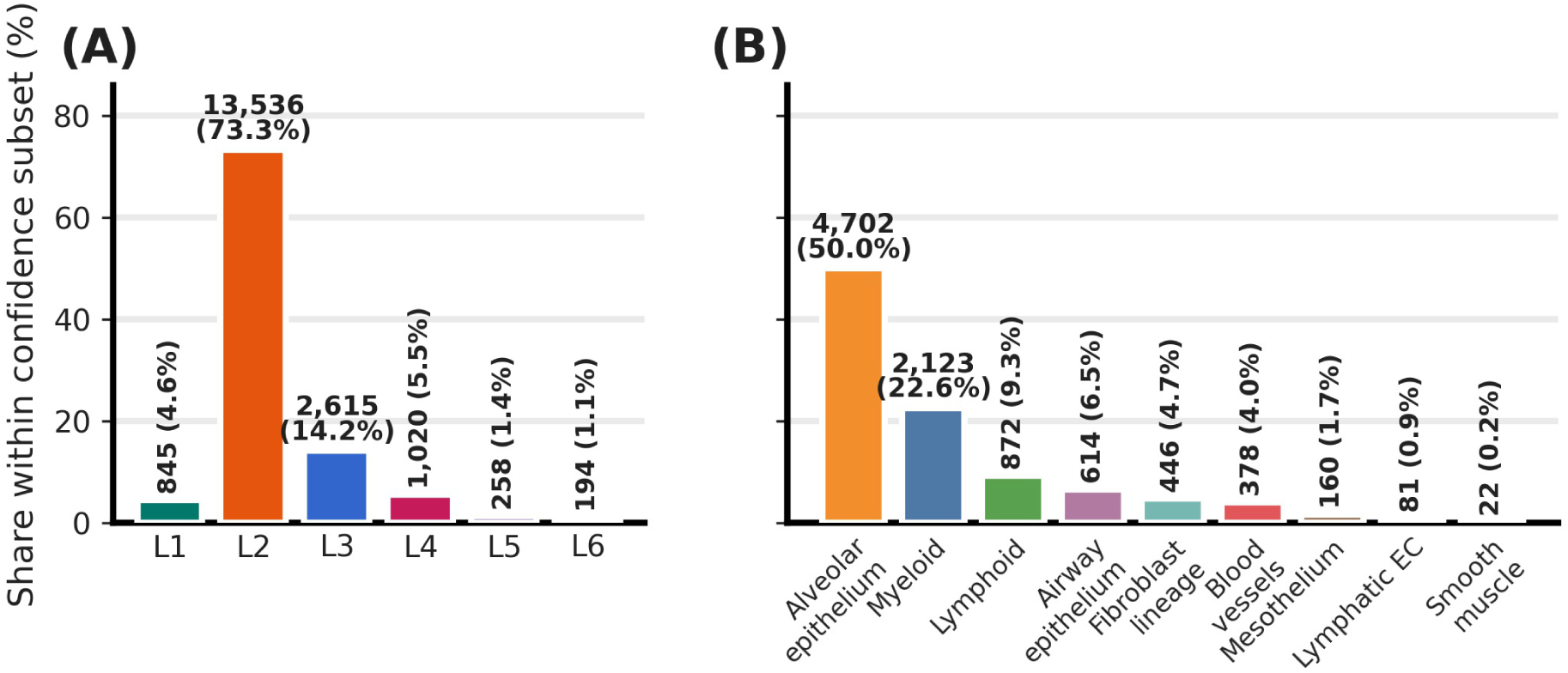
Composition of query cells from the original full-core HLCA Unknown set that were accepted by one Banovich-reference transfer but not the other. **(A)** Cells confidently placed by MESIC transfer but not by expression-space transfer (*n* = 18,468), split by transferred MESIC cluster. For context, in the Banovich reference subset, L2 was the main assigned cluster for blood-vessel, lymphatic-endothelial, fibroblast-lineage, and smooth-muscle cells, while also containing an alveolar epithelial group; an L2 transfer is therefore a cluster-level placement rather than a single level-2 cell-type label. **(B)** Cells confidently labeled by expression-space transfer but not by MESIC transfer (*n* = 9,398), split by transferred level-2 label. Bar labels give cell counts and percentages within each panel.

## 3 Materials and Methods

### 3.1 Semantic embeddings of gene summaries

We began with the 33,703 human gene symbols included in GenePT (Chen and Zou, 2025) and used NCBI Gene summaries collected in 2026 (Brown et al., 2015). Genes with an empty NCBI summary were excluded, leaving 21,788 genes. For each retained gene, the embedding input consisted only of its summary text after removing the gene-symbol label, terminal provider and source attributions, NCBI page-footer text, and NCBI Datasets download boilerplate.

We embedded each summary using the PubMedBERT model pretrained on PubMed abstracts and full text (Gu et al., 2021). Final-layer token representations were averaged to obtain one 768-dimensional vector per gene, producing a 21,788-by-768 gene-by-embedding matrix *M* . We mean-centered each embedding coordinate across genes and denote the centered matrix by *M̃*.

### 3.2 Construction and core-gene-based selection of MESIC components

We used Horn’s parallel analysis (Horn, 1965; Buja and Eyuboglu, 1992) to choose the number of semantic directions retained from *M̃*. Horn’s procedure compares observed PCA eigenvalues with eigenvalues expected after permuting the data, retaining directions whose variance exceeds this null expectation. This selected 50 directions, explaining 82.2% of the variance in *M̃*.

We then used a sparse PCA-like method to obtain a 768-by-50 semantic direction matrix *W* and the MESIC gene-embedding matrix *P* = *M̃W*, whose entry *P_gc_* denotes the value of gene *g* on component *c*. The method iteratively optimized an objective that combined a reconstruction loss with penalty terms that promoted near-orthogonality among the directions in *W* and concentration of each component’s large-magnitude entries in a small subset of genes. To select a final embedding, we obtained candidate *P* matrices from random initializations of *W* and a sweep over penalty weights.

For each fitted candidate, after this optimization was complete, we defined each MESIC component’s core-gene set as the fixed point of the following pruning iteration. Starting from the full vector for that component, we computed the participation ratio (PR), 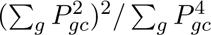, and rounded it to its nearest integer. We retained that number of genes with the largest *|P_gc_|*, set all other entries in the component vector to zero, and repeated this calculation on the pruned vector until the retained gene set no longer changed. This procedure converges because the number of retained genes is non-increasing and bounded below by one. The pruning step was not part of the optimization.

Among these candidates, we selected the final *P* using two component-level summaries, the median core-gene-set size and the median retained squared-coordinate fraction, defined as the sum of 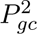 over core genes divided by the sum of 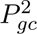 over all genes. This selection was completed before analyzing any expression dataset, cell label, disease label, or external biological annotation. After the final *P* was selected, downstream analyses used the unpruned *P*. Full details are provided in the Supplementary Methods.

### 3.3 Gene-type prediction benchmark

Gene-type labels were taken from GENCODE v50, corresponding to Ensembl release 116 (Frankish et al., 2023), and matched to modeled genes by exact gene symbol. Genes without a matched label and gene-type classes represented by fewer than 20 genes were excluded, leaving 21,674 genes across 13 classes. GENCODE labels were not used to construct or select *P*.

We evaluated two embedding matrices on the same retained genes, labels, and cross-validation splits: the 768-dimensional PubMedBERT embedding matrix *M* and the 50-dimensional MESIC gene embedding matrix *P*. Evaluation used five repeats of five-fold stratified cross-validation, giving 25 held-out folds. For each embedding matrix, we trained logistic-regression, random-forest, and XGBoost classifiers. Performance was summarized by accuracy, balanced accuracy, and macro-F1, where macro-F1 is the unweighted mean of the class-specific F1 scores. Benchmark results are reported in Table 1; label-processing rules, classifier settings, and additional implementation details are provided in the Supplementary Methods.

### 3.4 Annotation of MESIC component core genes

After the MESIC gene embedding *P* and component core-gene sets had been fixed, we annotated each component by testing its core genes for enrichment against external annotation terms, where each term was a Gene Ontology biological process, molecular function, or cellular component term, a Reactome pathway, or an RNAcentral HGNC-linked RNA gene class (Ashburner et al., 2000; The Gene Ontology Consortium, 2026; Ragueneau et al., 2026; RNAcentral Consortium, 2021). Annotation was used only for interpretation and did not alter *P*, component selection, cell projection, or downstream clustering.

Enrichment was tested with the hypergeometric upper tail using the 21,788 genes represented in *P* as the background. When computing the size of each external annotation term, we counted only genes that were also present in *P*, because term size determines the null expectation in the hypergeometric test. P-values were adjusted separately within each component using the Benjamini–Hochberg procedure (Benjamini and Hochberg, 1995). Terms with adjusted *q ≤* 0.05 and at least two overlapping core genes were retained as supported annotations.

For each component, the retained annotation terms were grouped into themes to reduce redundancy. Terms with largely overlapping core-gene support were placed in the same theme. The most significant term in each theme was used as the theme label, and the genes assigned to that theme were the union of core genes supporting its terms.

Compact component titles were derived from up to two leading themes. Components with no retained annotation terms were left unresolved. Full term-size filters, theme-grouping thresholds, ordering rules, and annotation-ledger details are provided in the Supplementary Methods.

### 3.5 Projection of single-cell expression profiles onto MESIC components

For each expression dataset, expression genes were matched to rows of *P* by exact gene symbol and kept in the same order in both matrices. The matched expression matrix was then projected onto the matched rows of *P* :

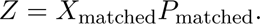

Rows of *Z* correspond to cells, and columns correspond to MESIC components. Dataset-specific analyses below state the expression source, matched-gene count, component exclusions, and standardization population.

### 3.6 Detecting and interpreting cardiomyocyte outliers in MESIC cell embeddings

We analyzed 15,846 cardiomyocytes from the human dilated and hypertrophic cardiomyopathy single-nucleus RNA-seq dataset of Chaffin et al. (Chaffin et al., 2022). The expression matrix *X* contained log-normalized CellBender-derived expression values, and 18,558 expression genes matched *P*. After projection, component values were centered and scaled across cardiomyocytes. Component C25 was excluded because none of its core genes overlapped the matched expression genes, leaving 49 components for downstream analysis.

For each cell, we computed the squared Mahalanobis distance from the cardiomyocyte population mean in the standardized 49-component embedding. The covariance matrix used in this distance was estimated with Ledoit–Wolf shrinkage (Ledoit and Wolf, 2004). We defined the top 1% of cells by squared Mahalanobis distance as outliers, yielding 159 cells, and tested enrichment of disease labels among outliers using one-sided Fisher exact tests.

We fit an elastic-net-penalized logistic-regression model (Zou and Hastie, 2005) to summarize which components distinguished the 159 outliers from the remaining 15,687 cardiomyocytes. The model used the 49 standardized component values as predictors. After selecting the elastic-net hyperparameters by cross-validation, we fit one final elastic-net logistic-regression model on all cardiomyocytes and used its coefficients for component interpretation. For component *j*, the squared-coefficient fraction was

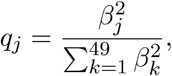

where *β_j_* is the final logistic-regression coefficient for component *j*. This fraction summarizes relative coefficient magnitude only; the sign of *β_j_* indicates whether higher or lower component values were associated with the outlier group. Hyperparameter selection and model-selection details are provided in the Supplementary Methods.

### 3.7 Unsupervised clustering of HLCA MESIC cell embeddings

We analyzed 121,894 cells from the Banovich/Kropski subset of the Human Lung Cell Atlas (HLCA) core atlas (Sikkema et al., 2023), referred to here as the Banovich subset. The expression matrix *X* contained log-normalized expression values, and 17,522 expression genes matched *P*. Component C25 was excluded because its core genes did not overlap the matched expression genes, leaving 49 components. After projection, component values were centered and scaled across Banovich-subset cells.

We constructed weighted nearest-neighbor graphs from the standardized 49-component cell embeddings using Euclidean distance. Leiden clustering (Traag et al., 2019) was run across neighborhood sizes and resolution values, with 1,000 random seeds for each setting. Settings were evaluated by recurrence of the same cluster count and by seed-to-seed agreement measured with adjusted Rand index (ARI) (Hubert and Arabie, 1985). The selected setting used 40 nearest neighbors and resolution 0.275; full stability diagnostics are provided in the Supplementary Methods.

At the selected setting, 983 of 1,000 Leiden runs produced six clusters. Consensus labels were assigned from these 983 six-cluster partitions. Partitions were aligned to a medoid partition, and each cell was assigned to the cluster receiving the most votes across aligned partitions. Consensus confidence was the fraction of aligned partitions voting for the assigned cluster. Cells with confidence at least 0.95 were assigned to one of six clusters, referred to as L1–L6, and cells below 0.95 were labeled uncertain.

HLCA annotations and UMAP coordinates were not used for graph construction, parameter selection, partition alignment, or consensus labeling. UMAP was used only for visualization.

As a robustness analysis, we also applied spectral clustering (Ng et al., 2002) using the same standardized 49-component cell embeddings and nearest-neighbor graph-construction procedure. We computed a spectral embedding from the graph, clustered the spectral coordinates with KMeans across 1,000 random seeds, and formed consensus labels using the same medoid-alignment and voting procedure used for Leiden. Full spectral-clustering parameters and stability diagnostics are provided in the Supplementary Methods.

### 3.8 Comparison of HLCA clusters with atlas annotation levels

After the MESIC consensus labels were fixed, we compared the six high-confidence clusters with HLCA annotation levels. Cells labeled uncertain by the consensus rule were excluded, leaving 117,928 cells. For the primary comparison, the same 117,928 cells were used at every annotation level, and entries without an assigned HLCA label at that level were treated as a separate unassigned category. For each annotation level, agreement with the MESIC clusters was summarized using adjusted Rand index (ARI), adjusted mutual information (AMI), and V-measure (Hubert and Arabie, 1985; Vinh et al., 2010; Rosenberg and Hirschberg, 2007). The annotation level with the highest ARI was reported as the HLCA resolution most closely matching the six-cluster MESIC partition. As a diagnostic for annotation levels with incomplete coverage, we also recomputed ARI after excluding cells without an assigned HLCA label at that level.

This comparison was used only to describe the biological scale of the unsupervised clusters. HLCA annotations were not used to construct the neighbor graphs, select the Leiden setting, align partitions, or assign consensus labels.

### 3.9 Multinomial logistic-regression summaries of HLCA MESIC clusters

We fit a six-class elastic-net multinomial logistic-regression model (Zou and Hastie, 2005) to summarize which components distinguished L1–L6 labels among cells with high-confidence consensus assignments. The model used the standardized 49-component cell embeddings as predictors and balanced class weights.

Regularization strength and the elastic-net mixing parameter were selected by stratified cross-validation, balancing multiclass balanced accuracy with coefficient sparsity. After selecting these hyperparameters, we fit one final elastic-net multinomial logistic-regression model on all high-confidence cells and used its coefficients for component interpretation. Hyperparameter grids, model-selection details, and held-out performance summaries are provided in the Supplementary Methods.

Because the predictors were the same component values used to define the clusters, heldout classification performance was not treated as independent validation. We used the fitted coefficients only as descriptive summaries. For Figure 6, squared-coefficient fractions were computed within each cluster as 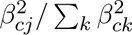, where the denominator sums over all 49 component predictors for that cluster.

### 3.10 Transfer of HLCA cells with Unknown level-2 labels

We used the 53,331 extended-atlas cells whose HLCA level-2 label (ann_level_2) was Unknown as query cells. The Banovich subset supplied the reference cells for both transfer analyses.

For the MESIC transfer, query cells were projected onto the same C25-excluded 49-component space used for Banovich-subset clustering and standardized with the Banovich-subset component means and standard deviations. Reference labels were the Banovich-subset consensus labels L1–L6, with uncertain consensus cells retained as an additional possible transfer outcome.

For the expression-space transfer, reference and query coordinates were taken from the 30-dimensional single-cell ANnotation using Variational Inference (scANVI) embedding provided by HLCA, and reference labels were the Banovich-subset level-2 labels.

Both transfers used distance-weighted 50-nearest-neighbor voting with Euclidean distance. For each query cell, normalized neighbor weights were summed within each reference label, and the label with the largest total weight was assigned. That largest label weight was used as the transfer confidence, and assignments with confidence at least 0.8 were considered confident. For the MESIC transfer, only confident assignments to L1–L6 were counted as reference-cluster placements; confident assignments to the uncertain reference label were recorded separately.

## 4 Discussion

This paper introduced MESIC and tested it in two stages, on the components themselves and on cell embeddings of two datasets. Compressing the PubMedBERT embeddings of 21,788 gene summaries into 50 components retained the gene-type information of the original embeddings. Forty-two components were enriched for Gene Ontology, Reactome, or RNAcentral themes. In cardiomyocytes, the cells farthest from the others in component space were enriched for HCM. Calcium handling, mitochondrial respiration, and fatty-acid metabolism were among the components that most distinguished the outliers, and each has been implicated in HCM (Coppini et al., 2013; Helms et al., 2016; Nollet et al., 2023). In the HLCA Banovich subset, unsupervised clusters of the cell embeddings agreed most closely with level-2 annotation, with several level-2 cell labels mapping predominantly to a single cluster despite large differences in label abundance. Several of the components that separated the clusters described the cells in them, with antigen-recognition genes for lymphoid cells, innate-defense genes for myeloid cells, and axonemal genes for ciliated airway epithelium. Used as a reference for label transfer, the clusters gave high-confidence cluster-level placements to about half of the extended-atlas cells that the original HLCA transfer had left without a level-2 label. Expression-space transfer from the same reference cells did not label most of those cells.

Across these analyses, the same components served outlier detection in heart, clustering in lung, and label transfer across the atlas without being refit, and in each case the result was characterized by named genes rather than by a search for markers after the fact. One lung result shows what a component label is and is not. C20 was named from sperm-flagellum annotations, but it separated the airway-enriched L5 cluster because its core genes encode the axonemal machinery that motile airway cilia share with sperm flagella (Brooks and Wallingford, 2014). The label is shorthand for an inspectable gene set, and that gene set carried meaning beyond the context that supplied the label.

MESIC rests on the assumption that a language model captures the functionally relevant content of a gene summary. C20 is that assumption working, and the results at the gene and cell levels support it in aggregate. Three tasks follow. First, the components were built from raw NCBI summary text, and some reflect curation templates and nomenclature in addition to biology. These template-driven components are where the assumption is weakest. C13, for example, groups biologically different genes whose summaries share Greek-letter nomenclature. Separating reusable biological language from database phrasing before embedding should sharpen the components. Second, the PCA-like method we use is one route to components that are sparse and traceable to genes. Comparing it with other interpretable factorizations, and rebuilding the components from newer biomedical and general-purpose embedding models, will show how much the component space depends on these choices. Third, the regression summaries reported here identify the components that distinguish a cell group. They do not explain individual cells. Outlier status was defined by joint distance across all components, so a cell can be outlying through an unusual combination of components even when no single component is extreme. Decomposing each cell’s distance into contributions from each component, accounting for their covariance, would extend the explanation from the outlier group to each outlying cell and remains for future work.

The components are fixed, so they give cell states a stable reference across time, treatment, and tissue. Longitudinal, perturbational, and spatially resolved data could be followed on the same components. The construction of the components is also not specific to genes. Any biological entity with curated text descriptions, including proteins, metabolites, microbial taxa, drugs, and phenotypes, could receive interpretable coordinates in the same way. Cells, perturbations, diseases, and candidate interventions would then share one semantic framework that connects an observed cell state to the experiments and therapeutic hypotheses relevant to it. In the near term, MESIC complements existing cell embeddings rather than replacing them, by supplying coordinates in which a result can be explained through genes.

## Supporting information

Supplemental Table 1. Component core genes

Supplemental Table 2. Component themes and gene summaries

## Author Contributions

X.D., M.A., and V.P. conceived the study, developed the methodology, designed the analyses, evaluated the results, and interpreted the findings. X.D. implemented the computational workflow, ran the code, curated the data, generated the figures, and wrote the initial manuscript draft. M.A. and V.P. critically revised the manuscript and provided intellectual input. V.P. supervised the project and acquired funding. All authors approved the submitted version and agree to be accountable for the content of the work.

## Acknowledgements

This research was supported by the Intramural Research Program of the National Institute of Diabetes and Digestive and Kidney Diseases (NIDDK) within the National Institutes of Health (NIH). The contributions of the NIH author(s) were made as part of their official duties as NIH federal employees, are in compliance with agency policy requirements, and are considered Works of the United States Government. However, the findings and conclusions presented in this paper are those of the author(s) and do not necessarily reflect the views of the NIH or the U.S. Department of Health and Human Services.

## Supplemental Data

The supplementary material accompanying this preprint includes a Supplementary Methods and Figures PDF and three supplemental data files.

The archive MESIC_supplemental_data_non_xlsx_20260904.zip contains the small frozen inputs required by the repository reproduction workflow. It includes the NCBI summary input files used for PubMedBERT embedding, the frozen gene-set annotation files used for component annotation, and the SCP1303 cardiomyocyte cell-ID file used to reconstruct the cardiomyocyte expression input. The archive also includes README files and SHA-256 checksum manifests.

The file selected_component_core_genes_wide.xlsx lists the fixed-point participation-ratio core genes for the 50 MESIC components. Columns C01–C50 correspond to MESIC components, and each column lists the genes retained in that component core.

The file component_card_theme_gene_summary_texts_grouped.xlsx lists the component-theme table used to inspect and label the component cards. It includes component identifiers, component display names, theme identifiers, theme names, grouped core-gene symbols, and the corresponding cleaned NCBI gene-summary texts, with a notes sheet defining the fields.

## Data Availability Statement

Publicly available datasets and resources were analyzed in this study. NCBI Gene summaries were obtained from NCBI Gene (Brown et al., 2015); GENCODE v50 gene annotations are available from GENCODE at https://ftp.ebi.ac.uk/pub/databases/gencode/Gencode_human/ release_50/gencode.v50.annotation.gtf.gz. Gene Ontology, Reactome, and RNAcentral annotation resources were used for component annotation (The Gene Ontology Consortium, 2026; Ragueneau et al., 2026; RNAcentral Consortium, 2021).

The cardiomyopathy single-nucleus RNA-seq data are available from the Broad Single Cell Portal study SCP1303. The Human Lung Cell Atlas v1.0 data are available from CELLxGENE and the Human Cell Atlas data portal. Reproducible code, source manifests, and small reproducibility files are available at https://github.com/nihcompmed/MESIC. Other details about data and reproducibility are explained in Supplementary Materials.

