## Supplementary Methods and Figures for "Mapping Gene Expression to an Interpretable Semantic Space"

### 1 Supplementary Methods

#### 1.1 Overview

MESIC maps gene-summary text into a lower-dimensional, interpretable semantic space and then uses that space to analyze gene classes and single-cell expression datasets. The analysis begins with NCBI Gene summary text, embeds each retained gene summary with PubMedBERT, selects the number of semantic dimensions by Horn parallel analysis, and fits a sparse projection of the PubMedBERT space. The resulting gene components are then summarized by their core genes, annotated by external gene-set resources, compared with the original dense PubMedBERT representation, and used as a projection basis for single-cell expression data.

The methods below describe the statistical and data-processing procedures. Full shell commands, file-system paths, checksums, database URLs, and scheduler details are provided in the repository README and step-specific READMEs rather than repeated here.

#### 1.2 Gene-summary corpus

NCBI Gene summaries are live text records that are revised over time, and NCBI does not provide a versioned release of summary text corresponding to a past collection date. MESIC therefore provides two gene-summary corpus options. The first option is the frozen July 7, 2026 corpus used for the reported analyses; this option supports exact row counts and downstream number matching. The second option is a current-source corpus built from the NCBI bulk summary files available at the time of rerun; this option lets readers generate their own current PubMedBERT input, but it is expected to differ from the frozen corpus.

Both options begin from the same 33,703 human gene-symbol universe. In the frozen option, genes without non-empty July 2026 NCBI summary text were excluded before text embedding, leaving 21,788 genes. The embedded text field contained summary text only; aliases and other metadata were not appended to the model input.

The same summary-text cleaning rule was applied before embedding. The procedure removed leading gene-symbol prefixes, trailing boilerplate, and terminal bracketed provider/source suffixes. PubMed citation parentheticals and non-terminal source text were retained. A small set of 43 retained rows contained NCBI Datasets download-package page text in the archived source and were repaired before embedding.

The current-source option was used to quantify how much a later NCBI download would differ. In the September 4, 2026 audit, the current download produced 21,819 genes with non-empty cleaned summaries. Of these, 21,729 overlapped the frozen 21,788-gene input; 90 genes appeared only in the current input and 59 frozen genes were absent from the current input. Among overlapping genes, 21,171 summary texts were identical and 558 differed. Expressed as proportions, the current download recovered 99.7% of the frozen genes, 99.6% of current genes were present in the frozen input, the gene-set overlap was 99.3%, and 97.4% of overlapping genes had identical cleaned text. Thus, the frozen corpus is required for exact reproduction of the reported numbers, whereas the current-source corpus supports a transparent from-current-NCBI rerun that should remain largely overlapping but not identical.

#### 1.3 PubMedBERT gene embeddings

Each retained NCBI summary was embedded with the PubMedBERT-base uncased model trained on PubMed abstracts and full text. The embedding used mean pooling over final hidden states after applying the attention mask. Texts were truncated or padded to a maximum length of 512 tokens. Historical inference used a batch size of 32 and stored the merged gene-by-feature matrix as 32-bit floating point values.

The resulting dense PubMedBERT matrix, denoted $M$, had 21,788 rows and 768 columns. Rows followed the frozen gene-summary input order. This matrix was not centered at the embedding stage; centering was applied only in downstream dimension-selection and projection analyses.

#### 1.4 Dimension selection

The number of semantic dimensions was selected by Horn parallel analysis. The PubMedBERT matrix $M$ was centered across genes to obtain $\widetilde{M}$. The real eigenvalue spectrum was computed from $\widetilde{M}$. Null spectra were generated by independently permuting each embedding coordinate across genes, preserving the marginal distribution of each coordinate while breaking gene-level covariance.

The analysis used 200 null permutations and compared the real eigenvalues with the 95th percentile of the null eigenvalue distribution. The selected dimension was the first crossing where the real eigenvalue was less than or equal to the corresponding 95th-percentile null eigenvalue. This rule selected 50 dimensions. The first 50 dimensions explained 82.2% of variance in the centered PubMedBERT matrix; 43 dimensions explained 80% and 99 dimensions explained 90%.

#### 1.5 Sparse semantic projection

This section describes a sparse PCA-like method used to obtain a sparse, interpretable projection from the centered PubMedBERT gene-summary matrix. Let $\widehat{\mu}$ be the gene-wise column mean, let $X_B = M_B - \mathbf{1}\widehat{\mu}^\top$ be a minibatch of centered gene embeddings, and let $W \in \mathbb{R}^{768\times 50}$ contain the learned component directions. The projected gene values are

$$Z_B = X_B W, \qquad \widehat{X}_B = Z_B W^\top.$$

The repository stores the transpose of $W$ as `W`, following the scikit-learn convention in which component directions are rows. The repository field `Z` is identical to $P$ in the reported basis export.

For minibatch $B$, the optimized loss was

$$\begin{aligned}\mathcal{L}_B(W) = {} & \frac{1}{\widehat{\sigma}^2_{\text{robust}}} \frac{1}{|B|p} \sum_{g\in B} \omega_g \left\| X_{g\cdot} - \widehat{X}_{g\cdot} \right\|_2^2 \\ & + \lambda_{\text{score}} L_{\text{PCA}} A_t \frac{1}{|B|k} \sum_{g\in B} \sum_{j=1}^{k} \log\left(1 + \frac{\widetilde{Z}_{gj}^2}{\epsilon_z^2}\right) \\ & + \lambda_{\text{ortho}} L_{\text{PCA}} \frac{1}{k^2} \left\| W_{\text{norm}}^\top W_{\text{norm}} - I_k \right\|_F^2 ,\end{aligned}$$

where $p = 768$, $k = 50$, and $W_{\text{norm}}$ denotes column-normalized $W$. The row weight was

$$\omega_g = \frac{\tanh(e_g)/e_g}{|B|^{-1}\sum_{h\in B}\tanh(e_h)/e_h}, \qquad e_g = \left(p^{-1}\left\|X_{g\cdot} - \widehat{X}_{g\cdot}\right\|_2^2 + 10^{-12}\right)^{1/2}.$$

The robust variance $\widehat{\sigma}^2_{\text{robust}}$ was the square of 1.4826 times the median nonzero coordinate-wise median absolute deviation, estimated from a 5% gene subsample. $L_{\text{PCA}}$ was the ordinary rank-50 PCA reconstruction loss under the same robust weighting and scaling; for the selected run, $L_{\text{PCA}} = 0.2451$. The direct loading-sparsity term present in the implementation had weight zero in the reported sweep, so the active sparsity term was the score-sparsity penalty on $Z_B$.

Projected scores were normalized within each minibatch before applying the score-sparsity penalty:

$$\widetilde{Z}_{gj} = \frac{Z_{gj}}{\left(\sum_{h\in B} Z_{hj}^2 + 10^{-12}\right)^{1/2}}.$$

No exponential moving-average score normalization was used. The score threshold $\epsilon_z$ was initialized to $1/\sqrt{10000} = 0.0100$. During the first 2500 steps it was held fixed while the sparsity weight $A_t$ was cosine annealed from 0 to 1. After that warmup, $\epsilon_z$ was updated every 500 steps by Otsu thresholding of $|\widetilde{Z}_{gj}|$, with a floor at $1/\sqrt{|B|}$.

Optimization used JAX with 64-bit floating point enabled. Input embeddings were loaded as 64-bit floating point values for fitting; the exported $P$, `Z`, `W`, and centering mean were stored as 32-bit floating point arrays. The optimizer was Optax AdamW with global gradient-norm clipping at 1.0 and weight decay $10^{-4}$. The peak learning rate was $5 \times 10^{-3}$: it warmed linearly from $5 \times 10^{-5}$ to $5 \times 10^{-3}$ over 1250 steps, then decayed by a cosine schedule to $5 \times 10^{-4}$ over the remaining 3750 scheduled steps. Minibatches contained up to 10,000 gene rows, with the final minibatch in each shuffled pass smaller.

Each fit used 5000 scheduled gradient steps followed by a continuation phase at constant learning rate $5 \times 10^{-4}$. During continuation, the mean total loss was checked every 100 steps. Training stopped after five consecutive checks without a relative loss improvement greater than 0.5%, or after 5000 additional continuation steps. The selected run stopped after 5500 total gradient steps.

Model selection was based on a sweep of sparse projection fits. Randomly initialized fits were evaluated across four sparsity penalties $\lambda_{\text{score}} \in \{0.01, 0.1, 0.5, 1.0\}$ and 100 random seeds per penalty. PCA-initialized fits at the same penalties and ordinary dense PCA were included as comparators. All sparse projection fits used $\lambda_{\text{ortho}} = 0.1$.

Each fitted basis was summarized after assigning a fixed-point participation-ratio core to each component. The two model-selection summaries were the median core size across components and the median retained squared-coordinate fraction across components. For component $j$, let $C_j$ denote its fixed-point core gene set. The retained squared-coordinate fraction was

$$E_j = \frac{\sum_{g\in C_j} P_{gj}^2}{\sum_{g=1}^{N} P_{gj}^2}.$$

Thus, $E_j$ is the fraction of component $j$'s squared $P$ entries carried by its core genes. It is not PCA explained variance and is not the fraction of genes retained.

Selection was restricted to randomly initialized sparse-projection fits within the displayed Figure 1 axis ranges: median core size 14.5–27.5 and retained squared-coordinate fraction 0.013–0.056. After normalizing both axes within these displayed ranges, the selected fit was the point closest to the upper-left corner, favoring smaller core size and larger retained squared-coordinate fraction.

The selected model used 50 components, $\lambda_{\text{score}} = 1.0$, $\lambda_{\text{ortho}} = 0.1$, and random seed 21. Its median fixed-point core size was 18 genes and its median retained squared-coordinate fraction was 0.0506. The median one-pass dense participation-ratio count was 2369 genes. The orthogonality diagnostic, measured as Frobenius error per component of the loading Gram matrix, was 0.0240.

### 1.6 Component core genes

The fixed-point participation-ratio procedure used to define component core genes is described in the main Methods. In the implementation, genes were ranked within each component by decreasing $|P_{gc}|$. Stable sorting was used, so exact ties retained the frozen 21,788-gene input order. The maximum number of fixed-point updates was 1,000, and all 50 components in the selected basis converged before reaching this limit.

For the selected basis, the procedure yielded 4,412 component-core gene assignments representing 4,165 unique genes. A total of 223 genes appeared in more than one component core, with a maximum of four component cores for any single gene. Component core sizes ranged from 6 to 1,821 genes, with median 18; the largest core belonged to component C25. The retained squared-coordinate fraction ranged from 0.0080 to 0.7863 across components, with median 0.0506.

`selected_component_core_genes_wide.xlsx` is provided as Supplementary Data and contains the resulting component core-gene lists, with one column per component and core genes listed in within-component rank order.

### 1.7 GENCODE gene-type prediction benchmark

GENCODE gene-type labels were taken from GENCODE v50, corresponding to Ensembl release 116, and matched to the 21,788 genes by exact matching between the gene symbol and the `gene_name` field of GENCODE gene records. Only GENCODE records with feature type `gene` were used. When multiple GENCODE gene records matched the same symbol, the most frequent `gene_type` was assigned, with `gene_biotype` used only if `gene_type` was absent. Ties were resolved lexicographically. Of the 21,788 genes, 21,754 had a non-missing GENCODE label. Classes represented by fewer than 20 genes were excluded, leaving 21,674 genes across 13 classes. No genes were removed by the final multiple-retained-label ambiguity rule.

The benchmark compared the saved 768-dimensional PubMedBERT embedding matrix with the 50-dimensional MESIC gene embedding $P$ on the same retained genes. Cross-validation used five repeats of five-fold stratified splitting, giving 25 held-out folds. The split generator used random seed 20260717. Identical fold assignments were used for the PubMedBERT and MESIC embeddings, and the exact assignments are provided with the accompanying repository.

Logistic regression was fitted after standardizing each feature using means and standard deviations estimated from the training fold. It used the `lbfgs` solver and `max_iter`=5000. Random forest used 500 trees and square-root feature sampling. XGBoost used the multiclass soft-probability objective, 500 trees, maximum depth 4, learning rate 0.05, row subsampling 0.9, column subsampling 0.9, multiclass log-loss evaluation, and the `hist` tree method. No class weighting or classifier hyperparameter tuning was applied.

Accuracy was the fraction of held-out genes assigned to the correct class. Balanced accuracy was the unweighted mean of the class-specific recalls. Macro-F1 was the unweighted mean of the class-specific F1 scores, with zero-division cases assigned an F1 score of zero. Multiclass log loss was also calculated when predicted probabilities were available, but the main table reports accuracy, balanced accuracy, and macro-F1.

### 1.8 Component annotation

Component annotation was performed after the MESIC component core-gene sets had been fixed. This step was used only to interpret fixed component cores and was not used to construct the MESIC basis, select candidate embeddings, project cells, or choose downstream clustering parameters.

We tested component core genes against frozen gene-set snapshots from Gene Ontology biological process, molecular function, and cellular component terms, Reactome pathways, and RNAcentral HGNC-linked RNA gene classes. Gene sets were stored as GMT files and represented by gene symbols. Before testing, each gene set was restricted to genes represented in $P$; term sizes and overlaps were computed after this restriction, with the 21,788 $P$ genes serving as the background. Terms were excluded if they contained fewer than 5 genes or more than 500 genes for Gene Ontology and Reactome, or more than 2500 genes for RNAcentral.

For each component, over-representation of core genes in each retained gene set was evaluated using the hypergeometric upper tail. Terms with fewer than two overlapping component core genes were not retained. P-values were adjusted separately within each component using the Benjamini–Hochberg procedure, and terms with component-wise adjusted $q \leq 0.05$ were considered supported annotations.

Supported terms were then collapsed into nonredundant annotation themes using their overlapping component core genes. Terms were processed in a deterministic order based on adjusted $q$-value, nominal $P$-value, overlap size, source, and term name. A term was assigned to an existing theme when its overlap with that theme met either a Jaccard threshold of 0.50 or an overlap-coefficient threshold of 0.70; otherwise, it started a new theme. The leading term in each theme defined the theme label, and the theme gene set was the union of component core genes supporting terms in that theme. Compact component titles were derived from up to two leading themes. Components without retained themes were marked unresolved.

All component core genes were retained in an annotation ledger. Core genes covered by supported themes were assigned to their corresponding themes, while core genes not covered by a supported theme were placed in an unresolved category for that component. Such unresolved genes or components can arise when the relevant genes are absent from the supplied annotation libraries, when available terms are outside the prespecified size range, when term overlaps with the component core are too small, or when tested overlaps do not pass the adjusted-$q$ threshold.

After filtering, 13,029 gene sets were tested, producing 1,981 enrichment rows and 118 collapsed annotation themes. Eight components remained unresolved by the retained gene-set themes. `component_card_theme_gene_summary_texts_grouped.xlsx` is provided as Supplementary Data. It contains the grouped component-theme gene symbols and corresponding NCBI gene-summary texts used to inspect the component annotation cards.

### 1.9 Projection of expression data into MESIC space

Single-cell expression matrices were projected into MESIC space by matching expression genes to MESIC basis genes and multiplying the matched expression matrix by the corresponding rows of the gene embedding matrix:

$$Z = X_{\text{matched}} P_{\text{matched}}.$$

For the cardiomyocyte and Banovich/Kropski reference analyses, projected component values were centered and scaled within the analyzed dataset. For the unknown-cell transfer analysis below, query cells were centered and scaled using the Banovich/Kropski reference means and standard deviations so that query and reference cells were represented on the same scale.

For the cardiomyopathy single-nucleus dataset, we used the SCP1303 processed log-normalized CellBender expression matrix, not the raw-count matrix. The expression input contained 15,846 cardiomyocytes and 36,601 genes; the cardiomyocytes were the frozen GenePT cardiomyocyte subset provided in Supplementary Data. Of the 36,601 genes, 18,558 matched the MESIC basis. All 50 MESIC components were retained for projection. Component 25 had no matched core-gene overlap in this dataset but was retained at the projection stage so that downstream analyses could decide whether to filter it.

For the Banovich/Kropski subset of the Human Lung Cell Atlas, we selected cells from the integrated HLCA core object using `dataset=Banovich_Kropski_2020` and used the processed log-normalized `X` matrix, not `raw/X`. Genes were matched by exact gene symbol from `var/feature_name`. The prepared reference expression matrix contained 121,894 cells and 17,522 matched genes. Component 25 had no matched core-gene overlap in this lung reference and was removed before graph construction, leaving 49 standardized MESIC dimensions for HLCA analyses.

### 1.10 Cardiomyocyte outlier analysis

Cardiomyocyte outlier analysis used the standardized cardiomyocyte MESIC component values after removing component 25, leaving 49 dimensions. A Ledoit–Wolf covariance model was fit to all 15,846 cardiomyocytes, and squared Mahalanobis distances were computed for each cell. The top 1% of cells by global Mahalanobis distance were called outliers, yielding 159 cells. After the outlier set was fixed, enrichment of outlier status was tested against the metadata columns `SubCluster`, `cell_type_leiden0.6`, `disease`, and `biosample_id` using one-sided Fisher exact tests.

An elastic-net logistic regression model was used to summarize which MESIC components distinguished the top 1% Mahalanobis outliers from the remaining cardiomyocytes. The response was the binary top-1% outlier indicator, and the predictors were the 49 standardized MESIC component values remaining after removing component 25. The model used scikit-learn logistic regression with the SAGA solver, elastic-net penalty, no class weighting, maximum 200,000 iterations, tolerance $10^{-4}$, and random state 21. Hyperparameters were evaluated by 5-fold stratified cross-validation with shuffled folds and random state 21. The $C$ grid was 0.001, 0.002, 0.003, 0.005, 0.0075, 0.01, 0.015, 0.02, 0.03, 0.05, 0.075, 0.1, 0.15, 0.2, 0.5, and 1.0; the $\ell_1$-ratio grid was 0.1, 0.3, 0.5, 0.7, 0.9, and 1.0.

Model selection was constrained to hyperparameter settings that converged in all five folds. For each validation fold, predicted probabilities were also converted to a prevalence-matched binary call by labeling as positive the same number of cells as the true validation-fold outlier count. A setting was eligible if its mean validation AUROC, average precision, prevalence-matched F1, and prevalence-matched accuracy were within 0.01, 0.03, 0.03, and 0.005, respectively, of the best converged value for that metric. Among eligible settings, the selected model minimized the mean coefficient-energy effective number,

$$\left(\sum_j q_j^2\right)^{-1}, \qquad q_j = \frac{\beta_j^2}{\sum_\ell \beta_\ell^2},$$

where $\beta_j$ is the logistic-regression coefficient for component $j$. Ties were broken by larger mean top-five coefficient-energy fraction, larger mean AUROC, and larger mean prevalence-matched F1.

The selected setting used regularization strength $C = 0.075$ and $\ell_1$ ratio 0.9. Its mean validation AUROC was 0.947, mean average precision was 0.445, and mean F1 at prevalence was 0.428. The final model had 30 nonzero coefficients. The Ledoit–Wolf shrinkage was $5.33 \times 10^{-4}$. Squared Mahalanobis distances had median 46.7, 95th percentile 83.5, 99th percentile 104.7, and maximum 246.4.

### 1.11 HLCA Leiden consensus clustering

The Banovich/Kropski HLCA reference was clustered in the 49-dimensional standardized MESIC space. For each candidate nearest-neighbor value, neighbors were computed with Euclidean distance after excluding each cell as its own neighbor. Graph edge weights used an exponential transform of neighbor distance scaled by the median retained neighbor distance for that graph. The directed neighbor graph was symmetrized by retaining the larger of the two directed edge weights. Leiden clustering used the weighted `RBConfigurationVertexPartition` objective in `leidenalg`. Clustering was evaluated over neighbor values 10, 15, 20, 30, and 40; resolution values 0.02, 0.04, 0.06, 0.08, 0.10, 0.125, 0.15, 0.175, 0.20, 0.225, 0.25, 0.275, 0.30, 0.35, 0.40, 0.45, 0.50, 0.60, 0.70, and 0.80; and random seeds 1–1000.

Candidate settings were first summarized by the distribution of cluster counts across seeds. A setting was treated as noncollapsed if its modal cluster count exceeded two and the median largest-cluster fraction was below 0.90. Recurrent noncollapsed settings were those whose modal cluster count appeared in more than 900 of 1000 seeds. Exact pairwise adjusted Rand index was then computed across seed-specific partitions for recurrent settings carried forward to stability evaluation. The selected setting used 40 neighbors and resolution 0.275. It produced exactly six clusters in 983 of 1000 seeds. Across all 1000 seed-specific partitions for this setting, the median pairwise ARI was 0.977, and 95.3% of pairwise comparisons had ARI at least 0.95. For consensus construction, we retained only the 983 exact-six partitions; this gave $\binom{983}{2} = 482{,}653$ pairwise comparisons within the consensus subset, with median pairwise ARI 0.978.

Final Leiden labels were built from the 983 exact-six partitions at the selected setting. The consensus procedure selected the medoid seed with the highest mean ARI to the other exact-six partitions, aligned each partition's cluster labels to the medoid by maximum-overlap assignment on the contingency table, and assigned each cell to the medoid-aligned cluster receiving the most votes. Cell confidence was the fraction of the 983 aligned partitions voting for the assigned cluster. The medoid seed was 493. The primary manuscript label set used a 0.95 confidence threshold, assigning 117,928 cells and leaving 3,966 cells uncertain. Final Leiden label counts were L1: 39,728; L2: 26,839; L3: 23,677; L4: 17,730; L5: 6,794; L6: 3,160; and uncertain: 3,966.

Agreement with existing HLCA annotation levels was evaluated only after the Leiden consensus labels were fixed; annotation labels were not used for graph construction, Leiden parameter selection, medoid selection, or voting. The 0.95-threshold final label table was joined to the HLCA metadata by zero-based cell index. Cells labeled uncertain were excluded. For each of `ann_level_1` through `ann_level_5`, paired label vectors were formed from the final Leiden labels and the corresponding HLCA annotation values on the same 117,928 confidently assigned cells. In the primary comparison, blank annotation entries were retained as an explicit category, so all annotation levels were evaluated on the same cell set; coverage was reported as the percentage of those cells with a nonblank annotation. Agreement was summarized with adjusted Rand index, adjusted mutual information, and V-measure using `sklearn.metrics`. Adjusted mutual information used the arithmetic averaging convention. V-measure was computed as the harmonic mean of homogeneity and completeness. As a diagnostic, the same three metrics were also recomputed after removing blank entries for the annotation level being evaluated.

### 1.12 Auxiliary spectral consensus

As a robustness analysis for the HLCA clustering result, we also generated an auxiliary spectral-consensus clustering from the same 49-dimensional standardized HLCA MESIC matrix and nearest-neighbor cache used for the Leiden analysis. This branch was used only for the Figure 5 spectral comparison; it was not used to define the primary Leiden labels or any downstream elastic-net or unknown-cell transfer analysis.

Spectral affinities were built from the nearest-neighbor connectivity graph for neighbor values 30 and 40. The directed connectivity graph was symmetrized, self-edges were added, and spectral embeddings were computed with 12 dimensions using the ARPACK eigensolver and random state 20260822. For each neighbor value, KMeans was run with $k = 6$ for random seeds 1–1000 after row-normalizing the first six spectral coordinates. KMeans used 10 initializations, a maximum of 300 iterations, and Lloyd's algorithm.

The Figure 5 spectral panel used the 30-neighbor, six-cluster, threshold-0.95 spectral consensus result after aligning spectral labels to the Leiden label order by maximum overlap. The consensus procedure paralleled the Leiden consensus procedure: partitions were aligned to a medoid partition, and cell confidence was defined as the fraction of aligned partitions voting for the assigned cluster. The displayed spectral consensus had medoid seed 135, median pairwise ARI 1.000, 5th-percentile pairwise ARI 1.000, and 95th-percentile pairwise ARI 1.000 after rounding to three decimals. Spectral label counts were S1: 36,081; S2: 30,325; S3: 23,402; S4: 17,974; S5: 7,301; S6: 6,784; and uncertain: 27.

#### 1.13 HLCA multiclass elastic-net interpretation

A multinomial elastic-net logistic regression model was fit to interpret the Leiden labels in MESIC space. Cells with uncertain Leiden labels were excluded, leaving 117,928 confidently assigned cells. The predictors were the 49-dimensional standardized MESIC component values, and the outcome labels were L1–L6.

The model used scikit-learn multinomial logistic regression with the SAGA solver, elastic-net penalty, balanced class weights, maximum 4000 iterations, tolerance $10^{-3}$, and random state 20260824. A stratified 20% held-out test set was reserved before hyperparameter selection. Hyperparameters were chosen by four-fold stratified cross-validation on the remaining 94,342 cells, using shuffled stratified folds with random state 20260824. The $C$ grid was $10^{-4}$, $2 \times 10^{-4}$, $3 \times 10^{-4}$, $5 \times 10^{-4}$, $7 \times 10^{-4}$, 0.001, 0.0015, 0.002, 0.003, 0.005, 0.007, 0.01, 0.015, 0.02, 0.03, 0.05, 0.07, and 0.1; the $\ell_1$-ratio grid was 0.9 and 1.0. Cross-validation tracked accuracy, balanced accuracy, macro F1, weighted F1, multiclass log loss, and the number of nonzero coefficients.

For interpretation, model selection prioritized sparsity among models with near-optimal balanced accuracy. First, the maximum mean cross-validated balanced accuracy was identified. Models within 0.5 percentage points of that value were then treated as practically equivalent. Among these models, the selected model had the fewest mean nonzero coefficients across cross-validation folds, using $|\beta| > 10^{-8}$ as the nonzero threshold. Ties were broken by higher mean balanced accuracy, smaller $C$, and then larger $\ell_1$ ratio. This criterion selected $C = 0.007$ and $\ell_1$ ratio 1.0. Held-out balanced accuracy was 98.5%, held-out macro F1 was 0.971, and the final model fit on all confidently assigned cells had 114 nonzero class-component coefficients. Because the Leiden labels were derived from the same MESIC component values used as predictors, held-out classification performance was treated as a model diagnostic rather than as an independent validation of the clusters.

For the Figure 6 component-weight display, each raw class-specific coefficient was squared and normalized within its L1–L6 label:

$$100 \times \beta_{cj}^2 / \sum_k \beta_{ck}^2,$$

where $\beta_{cj}$ is the coefficient for component $j$ in class $c$. The figure displayed the five largest component weights for each Leiden label.

#### 1.14 Unknown-cell label transfer

Unknown-cell transfer analyzed 53,331 cells from the full HLCA object satisfying `cell_type=unknown` and `ann_level_2=Unknown`. Two transfer spaces were compared. The first used the 49-dimensional MESIC projection and transferred the final Leiden consensus labels from the Banovich/Kropski reference. For this MESIC transfer, query cells were projected from the processed log-normalized HLCA `X` matrix by exact gene-symbol matching and were standardized using the Banovich/Kropski reference means and standard deviations. The query projection used 17,207 genes shared with the reference-matched gene set. The uncertain reference label was retained as a possible transfer outcome, but only confident assignments to L1–L6 were counted as MESIC cluster placements.

The second transfer used the 30-dimensional scANVI (single-cell variational inference) embedding `obsm/X_scanvi_emb` available in HLCA and transferred Banovich/Kropski level-2 annotation labels. The same 53,331 query cells and the same 121,894 Banovich/Kropski reference cells were used for both transfer analyses.

Both transfer analyses used Euclidean 50-nearest-neighbor voting from the Banovich/Kropski reference. Neighbor votes used the HLCA-style distance weighting implemented in the repository. For query cell $i$, let $d_{ij}$ be the Euclidean distance to reference neighbor $j$, and let $\sigma_i$ be the standard deviation of the 50 neighbor distances for that query cell. The unnormalized neighbor weights were

$$\tilde{w}_{ij} = \exp\left\{-\frac{d_{ij}}{(2/\sigma_i)^2}\right\},$$

and normalized weights were

$$w_{ij} = \frac{\tilde{w}_{ij}}{\sum_{\ell=1}^{50} \tilde{w}_{i\ell}}.$$

Rows with $\sigma_i \leq 10^{-12}$ used uniform weights. Transfer uncertainty was defined as one minus the top-label weighted vote; labels with uncertainty at most 0.2 were considered confident, equivalent to requiring top-label vote weight at least 80%.

The MESIC Leiden-label transfer was confident for 28,114 query cells, whereas the scANVI level-2 transfer was confident for 19,044 query cells. Both methods were confident for 9,646 cells. MESIC -only confidence occurred for 18,468 cells, scANVI-only confidence occurred for 9,398 cells, and neither method was confident for 15,819 cells.

### 2 Supplementary Figures

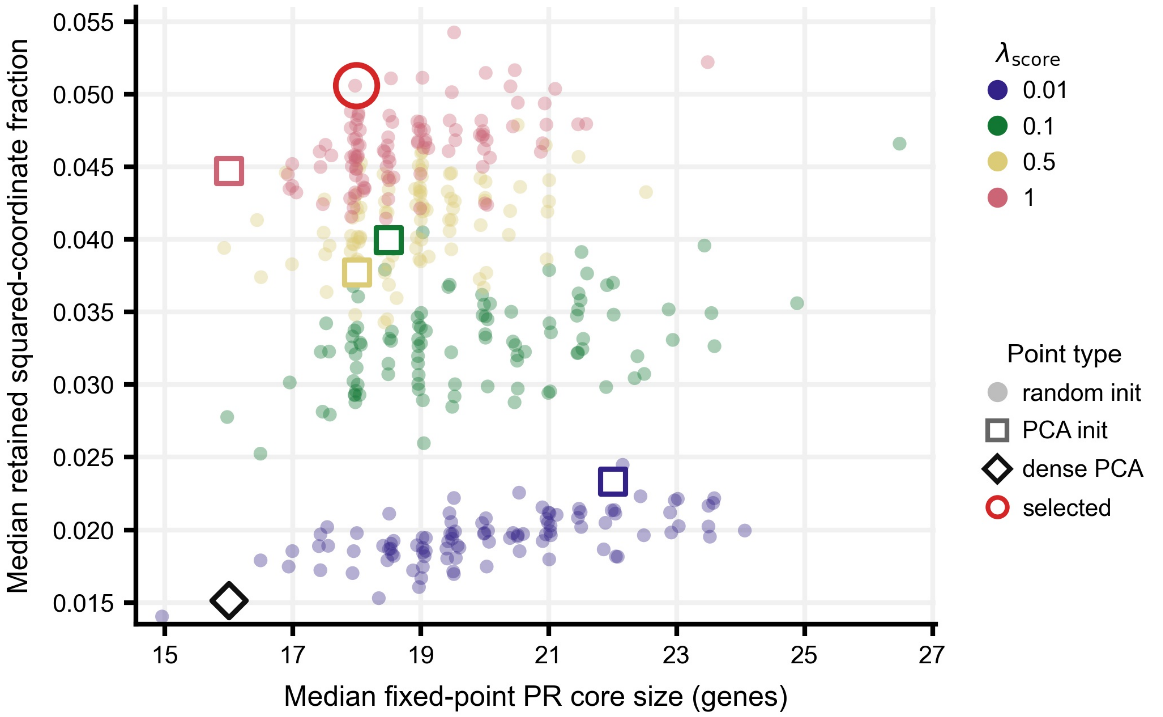


Figure 1: Selection of the MESIC gene embedding $P$. Each point represents one candidate embedding from the sparse projection model sweep. The x-axis shows the median fixed-point participation-ratio core size across the 50 components, and the y-axis shows the median retained squared-coordinate fraction across components. Colors indicate $\lambda_{\text{score}}$. Filled circles show randomly initialized runs, open squares show PCA-initialized runs, and the black diamond shows ordinary 50-component PCA. The selected model is the randomly initialized run with the smallest normalized distance from the upper-left corner within the fixed displayed model-selection ranges.
