## Supplementary material for "Mapping Gene Expression to an Interpretable Semantic Space": README.md

# NCBI Summary PubMedBERT Input

This package explains the provenance of the frozen NCBI gene-summary text table
used to generate the reported PubMedBERT embedding matrix.

The main reproduction path starts from file 03. Files 00--02 document how file
03 was produced.

## Files

```text
data/00_genept_released_ncbi_summary_snapshot_n33703.json.gz
```

GenePT-released NCBI summary snapshot downloaded from a GenePT-declared data
source. This file defines the 33,703-gene universe and provides the GenePT
baseline summary text used for comparison. It was not generated by the MESIC
pipeline.

```text
data/01_july2026_genept_vs_ncbi_gene_summary_snapshot_n33703.tsv.gz
```

Archived July 2026 comparison snapshot. It contains GenePT released text and
MESIC-collected July 2026 NCBI Gene-page text after collection, repair, and
rescue. It is not a raw NCBI release file.

```text
data/02_cleaned_genept_vs_july2026_ncbi_summary_comparison_n33703.tsv.gz
```

Cleaned 33,703-row comparison table generated from file 01. The cleaning step
removes leading `Gene Symbol <gene>` prefixes, trailing NCBI page boilerplate,
and terminal bracketed provider/source/date suffixes. It keeps PubMed citation
parentheticals and non-terminal provider/source text.

```text
data/03_pubmedbert_ncbi_summary_input_n21788.tsv.gz
```

Final frozen 21,788-gene PubMedBERT input. This is the text table used to
generate the reported embedding matrix. PubMedBERT embeds the
`embedding_text_summary_only` column.

## Processing Chain

File 00 is an external GenePT-released snapshot. The MESIC pipeline does not
recreate it from NCBI.

The July 2026 NCBI Gene-page collection scripts are archived under:

```text
pipeline/step00_ncbi_summary_sources/historical_20260707/
```

Those scripts document how the July 2026 NCBI text in file 01 was collected.
They are provenance scripts, not an exact future-rerun boundary, because NCBI
Gene pages and symbol mappings can change.

File 01 to file 02:

```bash
python pipeline/step00_ncbi_summary_sources/scripts/build_ncbi_summary_comparison.py
```

That repository script reads the same archived comparison snapshot through the
repository path:

```text
supplemental_data/ncbi_summary_pubmedbert_input/data/
  01_july2026_genept_vs_ncbi_gene_summary_snapshot_n33703.tsv.gz
```

File 02 to file 03:

```bash
python pipeline/step01_ncbi_summary_pubmedbert_input/scripts/build_frozen_ncbi_summary_pubmedbert_input.py
```

File 03 to PubMedBERT embeddings:

```bash
mkdir -p results/step01_ncbi_summary_pubmedbert_input
gzip -dc supplemental_data/ncbi_summary_pubmedbert_input/data/03_pubmedbert_ncbi_summary_input_n21788.tsv.gz \
  > results/step01_ncbi_summary_pubmedbert_input/n21788_geneSPAR-2026-current-summary-only_no-alias-rescue_citations-kept_external-download-cleaned.tsv
source pipeline/step02_pubmedbert_embedding/step02_env.sh
bash pipeline/step02_pubmedbert_embedding/submit_pubmedbert_embedding.sh
```

The Step00/Step01 folders also provide a current NCBI FTP route:

```bash
PYTHON_BIN=python bash pipeline/step00_ncbi_summary_sources/run_current_ncbi_ftp_summary_collection.sh
bash pipeline/step01_ncbi_summary_pubmedbert_input/run_current_ncbi_ftp_pubmedbert_input.sh
```

That command downloads current NCBI `gene_summary.gz` and
`Homo_sapiens.gene_info.gz`, maps the same 33,703 GenePT symbols by exact
official symbol, writes a Step00 current source table, and then writes a Step01
current PubMedBERT input table. It is a date-dependent audit, not an exact
regeneration of file 03. In the September 4, 2026 audit, the current FTP input
had 21,819 rows, 21,729 genes overlapped the frozen 21,788-row file 03, and
21,171 overlapping genes still had identical summary text.

## Row Counts

```text
00_genept_released_ncbi_summary_snapshot_n33703.json.gz: 33,703 genes
01_july2026_genept_vs_ncbi_gene_summary_snapshot_n33703.tsv.gz: 33,703 rows plus header
02_cleaned_genept_vs_july2026_ncbi_summary_comparison_n33703.tsv.gz: 33,703 rows plus header
03_pubmedbert_ncbi_summary_input_n21788.tsv.gz: 21,788 rows plus header
```

The final 21,788 PubMedBERT input rows are the July 2026 NCBI-summary rows
retained by Step01. The pipeline retained rows with non-empty July 2026 NCBI
summary text and excluded rows where the July 2026 NCBI text was empty. It also
repaired 43 rows contaminated by NCBI Datasets download-package UI text by
keeping only the NCBI title text before the `Homo sapiens` / `Gene ID` page
metadata.

## Checksums

Compressed file checksums are recorded in:

```text
checksums.sha256
```

Uncompressed SHA-256 checksums:

```text
3db721b7cfbc35795428c6a7ab4e8a9c59fb5f5ccb8c64af50dd3c97c2d4de2e  00_genept_released_ncbi_summary_snapshot_n33703.json
7e55ff9c6530bd2640fa79f1dfef8ea1c9b1dfec1dad6d4651441be29626758e  01_july2026_genept_vs_ncbi_gene_summary_snapshot_n33703.tsv
3e8b4a38da1b9a99c1004faf1fd21a48f7933afb25dde3a98985065bf59d908f  02_cleaned_genept_vs_july2026_ncbi_summary_comparison_n33703.tsv
65fc3b5b4a7158835d6a75c15f2f1b841668148393e90b06574cfc48034c854a  03_pubmedbert_ncbi_summary_input_n21788.tsv
```

## What Not To Claim

Do not describe file 03 as a raw NCBI download. It is a cleaned and selected
PubMedBERT input table.

Do not describe file 00 as generated by MESIC. It is a GenePT-released source
snapshot downloaded from a GenePT-declared data source.

Do not present a future live NCBI rerun as byte-identical reproduction of file
03 unless the resulting checksums match. Exact reproduction of the reported
downstream results starts from the frozen file 03.
