## Supplementary material for "Mapping Gene Expression to an Interpretable Semantic Space": README.md

# Supplemental Data

This folder contains reviewer-facing frozen intermediary files used for exact
reproduction of the reported MESIC analyses.

## NCBI Summary PubMedBERT Input

```text
ncbi_summary_pubmedbert_input/
```

This package documents the provenance of the frozen NCBI gene-summary text file
used as input to PubMedBERT. It contains the external GenePT-released summary
snapshot, the archived July 2026 GenePT-vs-NCBI comparison snapshot, the cleaned
comparison table, and the final 21,788-gene PubMedBERT input.

Start here:

```text
supplemental_data/ncbi_summary_pubmedbert_input/README.md
```

GENCODE v50 is not included here. Step07 uses the public versioned GENCODE v50
GTF download documented in `external_artifacts/README.md` and
`pipeline/step07_gene_class_prediction/README.md`.

## Step08 Annotation Gene Sets

```text
step08_annotation_gene_sets/
```

This package contains the frozen GO, Reactome, and RNAcentral GMT snapshots used
as Step08 enrichment inputs. These files are Supplemental Data because they are
not generated by Steps00-07 and exact Step08 reproduction should not depend on
future live database updates.

Start here:

```text
supplemental_data/step08_annotation_gene_sets/README.md
```

## Step09 Cardiomyocyte Sample Cell IDs

```text
step09_cardiomyocyte_sample/
```

This package contains the fixed 15,846-cell SCP1303 cardiomyocyte cell list
used to rebuild the Step09-ready cardiomyocyte expression artifact. The original
historical sample was described as a random subset but did not record the random
seed, so exact reproduction uses the supplied cell IDs rather than a new random
draw.

Start here:

```text
supplemental_data/step09_cardiomyocyte_sample/README.md
```
