## Supplementary material for "Mapping Gene Expression to an Interpretable Semantic Space": README.md

# Step08 Annotation Gene Sets

This Supplemental Data package contains the frozen gene-set GMT snapshots used
by Step08 annotation enrichment. These are not raw database downloads. They are
the exact GMT inputs consumed by `run_annotation_enrichment.sh`, included here
so the reported Step08 tables can be reproduced without depending on future GO,
Reactome, or RNAcentral updates.

## Files

```text
data/go/go_bp_human_from_gaf.gmt
data/go/go_cc_human_from_gaf.gmt
data/go/go_mf_human_from_gaf.gmt
data/reactome/ReactomePathways.gmt
data/rnacentral/rnacentral_hgnc_joined_rna_gene_sets.gmt
data/rnacentral/rnacentral_hgnc_joined_rna_gene_sets_audit.tsv
checksums.sha256
```

The five GMTs contain 14,399 input gene sets in total:

```text
GO biological process: 8,098 terms
GO cellular component: 1,244 terms
GO molecular function: 2,179 terms
Reactome pathways: 2,868 terms
RNAcentral RNA classes: 10 terms
```

## How Step08 Uses These Files

Step08 reads these GMTs as fixed enrichment libraries. The enrichment script
checks their SHA256 hashes before running. If any GMT has changed, the default
pipeline stops because the run is no longer an exact reproduction of the
reported MESIC annotation-enrichment result.

The RNAcentral audit TSV is not consumed by Step08. It is included only to make
the RNAcentral source-to-GMT construction inspectable.

Run Step08 from the repository root after Steps00-06:

```bash
bash pipeline/step08_annotation_enrichment/run_annotation_enrichment.sh
```

## GO Provenance

The GO GMTs were built from a human UniProt GAF and the matching GO OBO file
using:

```bash
python pipeline/step08_annotation_enrichment/scripts/convert_go_gaf_to_gmt.py \
  --gaf external_artifacts/go/HUMAN-uniprot.gaf.gz \
  --obo external_artifacts/go/go-basic.obo \
  --outdir supplemental_data/step08_annotation_gene_sets/data/go \
  --min-genes 5 \
  --max-genes 2000
```

The converter ignores `NOT` annotations, keeps IEA annotations by default,
propagates annotations to `is_a` and `part_of` ancestors, and writes separate
BP, CC, and MF GMT files.

Source metadata recorded from the files used to build the frozen GO GMTs:

```text
HUMAN-uniprot.gaf.gz
  gaf-version: 2.2
  generated-by: UniProt
  date-generated: 2026-06-18 08:16
  go-version: http://purl.obolibrary.org/obo/go/releases/2026-06-15/extensions/go-plus.ofn
  sha256: 258f6ea163375c036929478c400f25bf32e6d83df547cc4b864dfb4ec20c1ffb

go-basic.obo
  data-version: releases/2026-06-15
  sha256: c72fc198a86983d55e43aac585d1ffdbeb6e3601475b3f18b6045acdc0a0734c
```

The current GO human GAF path is:

```text
https://current.geneontology.org/annotations/gaf/HUMAN-uniprot.gaf.gz
```

For exact historical reproduction, use the frozen GMTs above. A future download
from a `current` URL may differ.

## Reactome Provenance

Reactome distributes `ReactomePathways.gmt` directly as a GMT inside
`ReactomePathways.gmt.zip`. The frozen file here is the unzipped GMT used by
Step08.

Source metadata recorded from the zip used to build the frozen Reactome GMT:

```text
ReactomePathways.gmt.zip
  source URL used historically: https://reactome.org/download/current/ReactomePathways.gmt.zip
  zip sha256: 8c1dbc8578431da5d2d5118262718c60b553a9be3398e93658daa069e4a9afd4
  zip member: ReactomePathways.gmt
  member size: 1,032,186 bytes
  member timestamp: 2026-06-21 06:23
```

The Reactome `current` download path is not immutable. Exact Step08
reproduction uses the frozen unzipped GMT in this package.

## RNAcentral Provenance

The RNAcentral GMT was built by joining RNAcentral
`id_mapping/database_mappings/` files through human HGNC mappings and retaining
only modeled Step06 background genes:

```bash
python pipeline/step08_annotation_enrichment/scripts/build_rnacentral_rna_gene_sets.py \
  --rnacentral-dir external_artifacts/rnacentral_database_mappings \
  --background results/step06_component_core_genes/selected_n21788_modeled_gene_background.txt \
  --outdir supplemental_data/step08_annotation_gene_sets/data/rnacentral
```

The builder requires these downloaded mapping files: `hgnc.tsv`, `mirbase.tsv`,
`mirgenedb.tsv`, `snodb.tsv`, `snopy.tsv`, and `snornadb.tsv`.

Source metadata recorded from the files used to build the frozen RNAcentral GMT:

```text
hgnc.tsv       rows: 9,024   sha256: 1ab93ceb6b257b29025616c4dee7b014c013597a1b2ed0e9f388efcdfa92f245
mirbase.tsv    rows: 87,467  sha256: 07680fad263300d87a557833930e1dc56492d6b3c74332be2810c63d306de496
mirgenedb.tsv  rows: 65,449  sha256: 8dcbcd952b6b294c1b90da91e4e245bbaae6d77f3924cc5d7f562fb258e33ee4
snodb.tsv      rows: 2,123   sha256: 431a77469c65f822c69ed1a9629186e7c04d9d269f75311239db4311f739ac93
snopy.tsv      rows: 2,634   sha256: ef63972e0fe40ce30e7c6001a522fbf901894b79bacbd87d318ce3f58e953840
snornadb.tsv   rows: 727     sha256: 3b68967c3dddf64579d13775336f687bdabd902fc68bf4799b5894e4c02ee093

generated audit TSV rows including header: 11
generated audit TSV sha256: 461c9025068030c75aa58d13a1d4d6b23d1205389041fd2e06b4cc4899681e3b
```

The RNAcentral source directory for current releases is:

```text
https://ftp.ebi.ac.uk/pub/databases/RNAcentral/current_release/id_mapping/database_mappings/
```

RNAcentral release outputs change over time. Exact Step08 reproduction uses the
frozen GMT in this package; rebuilding from a later `current_release` download
may produce a different GMT.
