## Supplementary material for "Mapping Gene Expression to an Interpretable Semantic Space": README.md

# Step09 Cardiomyocyte Sample Cell IDs

This Supplemental Data folder records the exact cardiomyocyte cells used when
preparing the Step09 cardiomyocyte expression artifact:

```text
external_artifacts/cardiomyocyte/sample_heart_data.h5ad
```

## File

```text
data/genept_cardiomyocyte_sample_heart_cell_ids_n15846.tsv
```

The table has two columns:

```text
sample_rank_0based
cell_id
```

It contains 15,846 unique SCP1303 cell IDs in the same order as the historical
GenePT `sample_heart_data.h5ad` object used for the existing MESIC results.
The original GenePT README described this H5AD as a random 10 percent
cardiomyocyte subset of the Broad Single Cell Portal SCP1303 cardiomyopathy
data, but it did not record the random seed. Therefore this release records the
selected cell IDs explicitly instead of asking reviewers to reproduce an
underspecified random sample.

## Use In The Pipeline

The Step09 preparation script consumes this cell list together with public
SCP1303 source files:

```bash
python pipeline/step09_expression_projection/scripts/prepare_cardiomyocyte_h5ad_from_scp1303.py
```

The script streams the SCP1303 log-normalized CellBender MatrixMarket file and
keeps only these 15,846 cells, preserving the row order in this TSV. It writes a
Step09-ready H5AD at:

```text
external_artifacts/cardiomyocyte/sample_heart_data.h5ad
```

This procedure is intended to reproduce the expression input needed by Step09
and downstream cardiomyocyte summaries. It is not a claim of byte-identical
reconstruction of the historical local H5AD because nonessential Scanpy objects
such as PCA, neighbor graphs, highly variable gene statistics, and the raw-count
layer are not reconstructed.
